# Longitudinal characterization of neuroanatomical changes in the Fischer 344 rat brain during normal aging and between sexes

**DOI:** 10.1101/2021.04.12.439510

**Authors:** Caitlin Fowler, Dana Goerzen, Dan Madularu, Gabriel A. Devenyi, M. Mallar Chakravarty, Jamie Near

**Affiliations:** Department of Biological and Biomedical Engineering, McGill University, Montreal, Canada; Centre d’Imagerie Cérébrale, Douglas Mental Health University Institute, Montreal, Canada; Center for Translational NeuroImaging, Northeastern University, Boston, MA, USA; Department of Psychiatry, McGill University, Montreal, Canada

**Author notes:** **Corresponding authors at:** CIC Pavillion, office GH-2113, Douglas Mental Health University Institute, 6875 Boulevard LaSalle, Montreal, Canada, H4H 1R3.

**Keywords:** Normative aging, magnetic resonance imaging, regional volumetry, longitudinal study, sex differences

## Abstract

Animal models are widely used to study the pathophysiology of disease and to evaluate the efficacy of novel interventions, crucial steps towards improving disease outcomes in humans. The Fischer 344 (F344) wildtype rat is a common experimental background strain for transgenic models of disease and is also one of the most frequently used models in aging research. Despite frequency of use, characterization of neuroanatomical change with age has not been performed in the F344 rat. To this end, we present a comprehensive longitudinal examination of morphometric change in 73 brain regions and at a voxel-wise level during normative aging in a mixed-sex cohort of F344 rats. We identified age- and sex-related changes in regions such as the cortex, hippocampus, cingulum, caudoputamen, and nucleus accumbens, which are implicated in memory and motor control circuits frequently affected by aging and neurodegenerative disease. These findings provide a baseline for neuroanatomical changes associated with aging in male and female F344 rats, to which data from transgenic models or other background strains can be compared.

**HIGHLIGHTS:**

- *In vivo* magnetic resonance imaging reveals altered neuroanatomy in aging Fischer rats
- Linear and curvilinear age effects exist in both grey and white matter structures
- Sex differences are primarily seen in grey matter structures
- This study clarifies normal aging trajectories across 73 brain regions in both sexes
- Improved understanding of normal aging will inform future pathological aging studies

## 1. INTRODUCTION

Aging is the predominant risk factor for the majority of diseases that reduce quality of life and shorten the lifespan, such as neurodegenerative diseases (Hou et al. 2019; Franceschi et al. 2018). Aging and age-related pathologies such as Alzheimer’s disease (AD) share many basic molecular and cellular processes including mitochondrial dysfunction, oxidative stress, cellular senescence, and inflammation, to which the brain is particularly susceptible (Franceschi et al. 2018; Farooqui and Farooqui 2009). Given these and other similarities, disentangling pathology from normal aging can be challenging, particularly in the early phases of disease when features such as cognitive impairment may be subclinical.

Preclinical studies in animal models represent an important step towards improving disease outcomes in humans. Importantly, studying aging in wildtype rodents provides a means of separating age-related changes from those arising due to pathology, given that disease phenotypes in rodents are generally introduced by means of transgene insertion or genetic knockout. A commonly used background strain for the development of transgenic lines is the Fischer 344 (F344) rat, which was recently used to generate a rat model of Alzheimer’s disease that spontaneously develops tau pathology (Cohen et al. 2013). The F344 rat is also one of the most frequently used strains for aging research (Gallagher, Stocker, and Koh 2011). Age-related changes in recognition and spatial memory (Marrone, Satvat, and Patel 2018; Guidi et al. 2014), hippocampal neurogenesis (G. A. Shetty, Hattiangady, and Shetty 2013) and inflammatory response (Mawhinney et al. 2011), and brain tissue metabolism (Harris et al. 2014; Fowler et al. 2020) have all previously been explored in the F344 rat. However, characterization of age-related change in brain structure in this rat strain has yet to be performed.

Brain volume measurements obtained through magnetic resonance imaging (MRI) techniques have been widely used to study aging in humans (for reviews, see (Fjell and Walhovd 2010; Hedman et al. 2012; Raz et al. 2005)), with somewhat fewer studies in rodents to date (Maheswaran et al. 2009; Driscoll et al. 2006; Gaser et al. 2012). The non-invasive nature of MRI makes it a unique tool for detecting and monitoring altered brain structure *in vivo*. Additionally, recent advances in MRI co-registration techniques have permitted the development of several widely used processing and analysis pipelines (Friedel et al. 2014; Tustison et al. 2014; M. Jenkinson et al. 2012) for longitudinal quantification of neuroanatomical change, with demonstrated success in preclinical studies (Rollins et al. 2019; Kong et al. 2018). All of these features, in combination with the significantly shorter lifespan of rodents compared to humans (approximately 21-26 months for the F344 rat (Chesky and Rockstein 1976)), provide a convenient and powerful means to study longitudinal neurobiological changes associated with normal aging across the lifespan.

Previous aging studies examining neuroanatomy in wildtype rodents are limited in number and inconsistent in how data are analyzed, with most studies reporting change in absolute brain volumes with age (von Kienlin et al. 2005; Oberg et al. 2008; Gaser et al. 2012; Hamezah et al. 2017; Casas et al. 2018) and only a few reporting change in relative brain volumes (accounting for total brain volume or intracranial volume), either at a regional level (Maheswaran et al. 2009; Driscoll et al. 2006) or at the level of individual voxels (Alexander et al. 2020). Furthermore, many of these studies are cross-sectional in nature or conducted in a single sex, reducing power and applicability. Additional studies are needed to establish the baseline for normal neuroanatomical change with age in the rodent brain, a necessary step towards understanding structural alterations in the context of pathology using transgenic models.

Studies that examine the influence of sex on brain changes during normal aging are an equally important part of establishing a baseline to which pathology-related neuroanatomical change can be compared. Significant differences between sexes exist regarding the risk for, and presentation and treatment of, age-related neurodegenerative diseases due to the underlying influence of sex chromosomes and hormones at the structural, functional, and biochemical level of the brain (Mazure and Swendsen 2016; Cosgrove, Mazure, and Staley 2007). Previous studies on the influence of sex on neuroanatomy have been performed cross-sectionally (Spring, Lerch, and Henkelman 2007) or relatively early in the lifespan (Qiu et al. 2018; Sumiyoshi, Nonaka, and Kawashima 2017; Kong et al. 2018; Corre et al. 2016), but to our knowledge, preclinical MRI studies examining the interaction between sex and age on neuroanatomy late into the lifespan do not currently exist. To this end, the present study describes a longitudinal analysis of brain morphometric change in both male and female F344 rats over the majority of the lifespan. Morphometric changes are reported both at the voxelwise level, and at the regional level in 73 unique brain regions. This work provides new insight into neuroanatomical trajectories associated with normal aging.

## 2. METHODS

### 2.1 Animals and study design

Homozygous Fischer 344/NHsd wildtype (WT) male and female rats were obtained from Envigo Laboratories (Madison, WI, United States; order code: 010) and bred within the Animal Care Facility at the Douglas Hospital Research Centre. Rats were weaned on postnatal day 21 and housed in pairs on a 12 hour light-dark cycle with *ad libitum* access to food and water. Both male and female experimenters handled and tested the rats, while animal staff caring for the rats were primarily female. All animal procedures and experiments were performed in accordance with the guidelines of the McGill University Animal Care Committee and the Douglas Hospital Research Centre Animal Care Committee.

MRI scans were acquired longitudinally at 4-, 10-, 16-, and 20-months of age, covering the majority of the adult rat lifespan. A total of 27 rats, (12M, 15F), were included in the study. 9 of 27 rats were scanned at only 4- and 10-months due to participation in a separate treatment study thereafter, leaving 18 rats (7M, 11F) to be studied at all four time points. Of these 18, 1 female died prior to the 16-month time point, and 4 males died prior to the 20-month time point, all of natural causes. Finally, one male and one female scan at 10-months and two female scans at 16-months were discarded after failing quality control. As such, the final number of scans for the four time points were 27 (12M, 15F), 25 (11M, 14F), 14 (7M, 7F), and 11 (3M, 8F), respectively (**Supplementary Table 1**). A linear mixed effects model (LME) was used to handle the imbalance in the number of scans per time point, as LMEs appropriately handle missing values in longitudinal analyses (Bernal-Rusiel et al. 2013).

### 2.2 MRI data acquisition

MRI data were acquired by C.F.F and D.M. at the Douglas Centre d’Imagerie Cérébrale using a 7 Tesla Bruker Biospec 70/30 scanner (Bruker, Billerica, MA, United States) with an 86 mm (diameter) volumetric birdcage coil for transmission and a four-channel surface array coil for signal reception (Bruker). The level of anesthesia (1-4% isoflurane in oxygen gas) was adjusted to maintain a breathing rate between 50-70 breaths per minute throughout the procedure and warm air (37 °C) was blown into the bore of the scanner to maintain a constant body temperature (SA Instruments, Inc., monitoring system, Stony Brook, NY, United States).

High-resolution 3D anatomical MR images were acquired using Rapid Acquisition with Relaxation Enhancement (RARE) using the following scan parameters: TR = 325 ms, echo spacing = 10.8 ms, RARE factor = 6, effective echo time = 32.4 ms, Field of View = 20.6 × 17.9 × 29.3 mm, matrix size = 256 × 180 × 157, slice thickness = 17.9 mm (along the dorsal/ventral direction), readout along the rostral/caudal direction, spatial resolution = 114 μm isotropic, 19m35s acquisition time. Following the scan, animals were allowed to recover from anesthesia and returned to group housing.

### 2.3 MRI pre-processing and registration pipelines

All images were processed in MINC 2.0 format. Preprocessing was performed using minc-toolkit-v2 (Vincent et al. 2016) and the MINC toolkit extras package (https://github.com/CoBrALab/minc-toolkit-extras), and the two-level model build Pydpiper module (Friedel et al. 2014) was used to co-register the pre-processed images into a common space. First, an in-house rat MRI preprocessing script within the minc-toolkit-extras package developed by G.A.D. (https://github.com/CoBrALab/minc-toolkit-extras/blob/master/rat-preprocessing-v3.sh) was used to perform the following sequential preprocessing steps: dimension reordering to standard MINC 2.0 ordering, image centring, whole image N4 bias field correction (Sled and Pike 1998; Tustison et al. 2010), individual foreground mask generation using the Otsu method (Otsu 1979) additional N4 bias field correction using the previously generated mask, affine registration to a Fischer 344 template average image, and a final N4 bias field correction using a template mask in native space. After pre-processing, images were quality controlled by D.G, during which images were visualised using the Display program in minc-toolkit-v2 and examined in each of the coronal, sagittal, and axial dimensions for motion artefacts, Gibbs ringing artefacts, proper bias field correction, and other image anomalies. Four scans were flagged and excluded from further analysis.

The remaining 77 scans which passed quality control were then co-registered using the two-level model build pipeline in Pydpiper (Friedel et al. 2014). In brief, subject-specific starting averages are created by rigidly registering scans at different time points to the Fischer 344 atlas template, followed by averaging. Iterative affine and non-linear registration and averaging is then repeated to produce an unbiased subject average. Subsequently, each subject specific average is rigidly aligned to the Fischer 344 atlas template space and the process is repeated to create an unbiased population average. This process creates deformation fields for each subject at each time point. The deformation fields can then be used to estimate the Jacobian determinant at each voxel, which reflects the amount of expansion or compression required to deform each individual anatomical image to the subject average (Chung et al. 2001). Deformation fields are then resampled into the common study space allowing comparison between subjects. This registration process generates two sets of Jacobian determinants which were used for subsequent structural analysis. The absolute Jacobian, composed of the sum of the linear plus the non-linear mappings, reflects the global changes in voxel volume. The relative Jacobian, composed of the non-linear mapping with residual affine components estimated and removed, reflects local, or relative changes in voxel volume. The Jacobian deformation fields were then blurred with a 400 micron full width half maximum Gaussian kernel to satisfy assumptions of normality required by the statistical models used to analyze the data.

### 2.4 Regional volume estimation

Using a Fischer 344 rat atlas generated from a 4-month cohort of wildtype Fischer 344 rats (Goerzen et al. 2020) resampled into the study common space, the volumes of 73 unique regions (120 when split across hemispheres, e.g. left and right Caudoputamen) were estimated using the *anatGetAll* function in RMINC_1.5.2.3 (J. Lerch et al. 2017). This function computes the volume of a region by counting the number of voxels with a given label and multiplying the Jacobian with the voxel volume at each voxel. Absolute and brain-size-corrected volumes (mm^3^) were generated from the absolute and relative Jacobians, respectively. Of the 120 delineated regions, 76 were classified as GM, 40 as WM, and 4 as CSF. Volumes aggregated within tissue types (grey matter, GM; white matter, WM; cerebrospinal-fluid-filled volumes, CSF) were also calculated to provide an overview of tissue-specific changes across the brain.

### 2.5 Statistical Analysis

Longitudinal changes in a) relative Jacobians at each voxel within the brain, b) absolute and c) brain-size corrected volumes for 73 regions, and d) volumes aggregated within tissue types, were modelled using linear mixed-effects (LME) modeling in R. (version 3.6.3 (R Core Team 2020); attached base packages: stats, graphics, grDevices, utils, datasets, methods, base; other attached packages: effects_4.4-4 (Fox and Weisberg 2019), RColorBrewer_1.1-2(Neuwirth 2014), readxl_1.3.1(Wickham and Bryan 2019), lmerTest_3.1-0(Kuznetsova, Brockhoff, and Christensen 2017), lme4_1.1-23 (Bates et al. 2015), tidyverse_1.3.0(Wickham et al. 2019), RMINC_1.5.2.3). LME models were used as they appropriately model the covariance structure resulting from repeated measurements in the same subjects and handle data with missing values (Bernal-Rusiel et al. 2013). In the mixed effects model, volumes were predicted by an age by sex interaction with a random intercept for each subject. Age was modelled using a quadratic function (poly(age,2)) to account for the possibility of non-linear changes with age, as has previously been demonstrated (Pfefferbaum et al. 2013; Kong et al. 2018; Tullo et al. 2019). To further justify this choice, Akaike information criterion (AIC; (Akaike, Petrov, and Csaki 1973)) was used to test if volume and voxelwise data were better fit using a linear or quadratic age term, using a Δ_i_ (AIC_i_-AIC_min_) threshold of 4, where AIC_min_ is the minimum of the *R* (number of models being compared) different AIC_i_ values (the minimum is at i=min) (Burnham and Anderson 2004). The larger the Δ_i_, the less likely the fitted model *i* is the best approximating model. Of 120 structures, 65 were best fit by a quadratic model, with 39 of those 65 reaching the Δ_i_, threshold of 4, indicating substantially less support for the model containing solely linear age term (**Supplementary Table 2**). Similarly, the majority of voxels demonstrated better fits using a quadratic age term. Thus, for consistency and comparability, all structures and voxelwise data were modelled using a quadratic age term.

The linear and quadratic components of the age term are henceforth abbreviated as poly(age,2)1 and poly(age,2)2, respectively. Main effects of sex (sexF) were evaluated as a group effect of females relative to males, while the interaction between age and sex (poly(age,2)1:sexF and poly(age,2)2:sexF) was also examined, again with males as the reference group. All continuous variables were z-scored so that coefficients (betas (β), or effect sizes) for each fixed effect would be standardized. As such, the standard betas indicate the number of standard deviations a regional volume has changed per standard deviation increase in the predictor variable. The False Discovery Rate (FDR) method (Benjamini and Hochberg 1995) was used to control the family-wise type I error at a level of 5% for each predictor. A fixed effect was considered significant when the FDR-corrected p-value (q-value) was < 0.05.

## 3. RESULTS

### 3.1 Absolute volumes increase with age and are larger in male Fischer rats

Linear mixed effects model results for absolute volumes aggregated within GM and WM tissues, as well as ventricular compartments containing CSF are visualized in **Figure 1A** and a full LME summary is shown in **Supplementary Table 3.** Briefly, aggregated absolute volumes all increased and demonstrated negative curvilinear relationships (downward-facing curve) with age (GM: poly(age,2)1 β=5.075, q=6.30E-25; poly(age,2)2 β=-1.258, q=5.53E-06; WM: poly(age,2)1 β=6.647, q=6.38E-29; poly(age,2)2 β=-1.159, q=5.36E-05; CSF: poly(age,2)1 β=6.439, q=5.31E-16; poly(age,2)2 β=-1.625, q=4.96E-03), were smaller in females (GM sexF β=-1.609, q=2.10E-10; WM sexF β=-1.372, q=3.07E-10; CSF sexF β=-1.325, q=1.04E-08), and demonstrated a negative linear age by sex interaction whereby volumes increased more sharply wth age in males than females (GM: poly(age,2)1:sexF β=-1.812, q=5.33E-07; WM poly(age,2)1:sexF β=-1.603, q=1.36E-05; CSF poly(age,2)1:sexF β=-1.647, q=2.28E-02).

**Figure 1:**
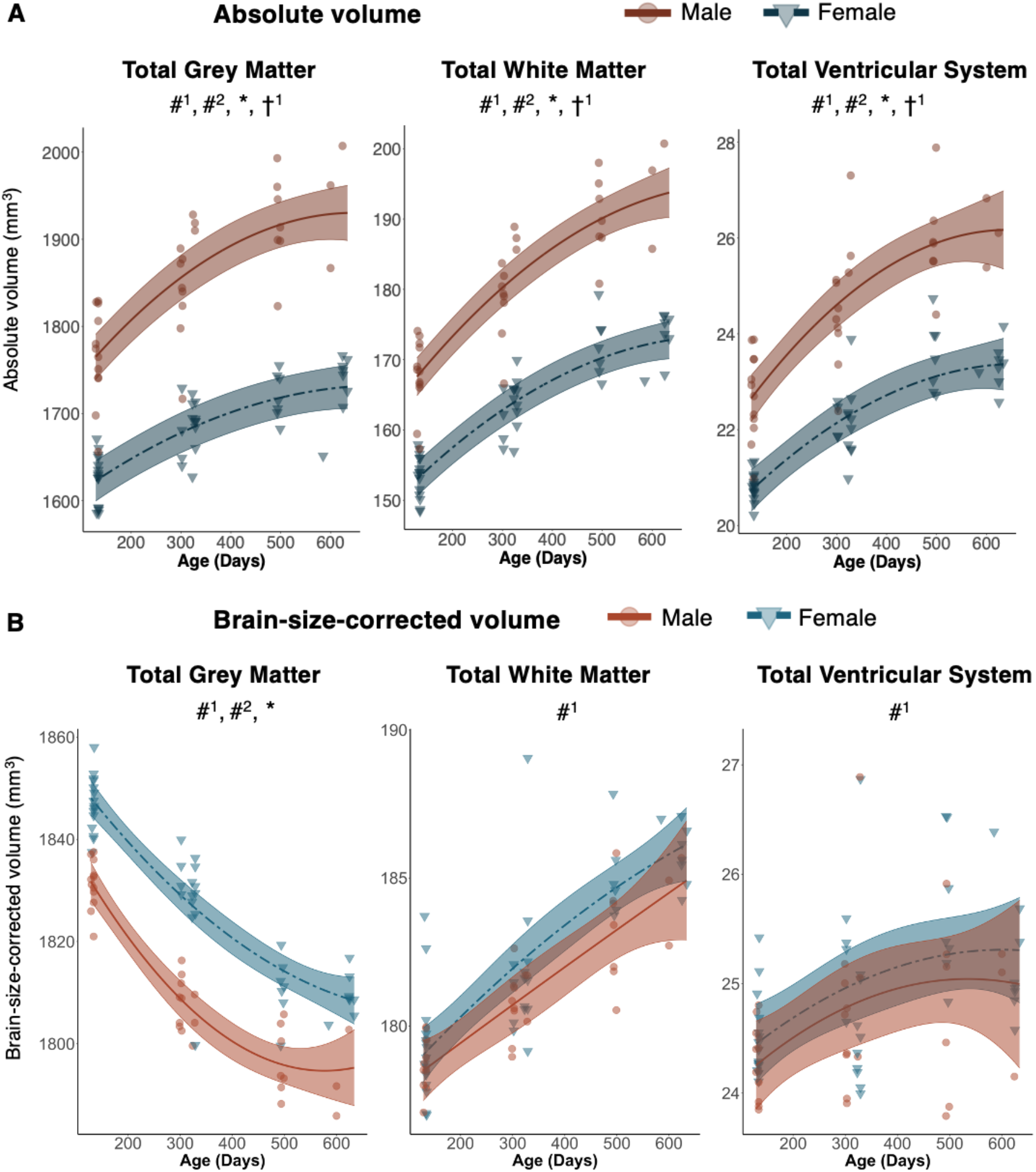
Absolute volumes summed across tissue type increase with age and are larger in males, while brainsize corrected volumes reveal different age and sex-dependent trends across tissue types. MRI data were acquired longitudinally in rats aged 4-, 10-, 16- and 20-months old. Main effects of age and sex, as well as age by sex interactions, were determined using linear mixed effects modelling. Each data point represents a single rat. The linear mixed effects model used to fit the data is represented by a line of best fit and 95% interval (shaded), split by sex. Data corresponding to males is shown using red circles with a solid line of best fit, while females are shown using blue triangles and a dashed line of best fit. Significance symbols are shown for each term in the model, where #^1^ and #^2^ represent the linear and quadratic age terms, * denotes a main effect of sex, and †^1^ and †^2^ represent age by sex interactions with a linear and quadratic age term, respectively. Multiple comparisons were corrected for using a 5% false discovery rate. Significance was determined by q<0.05.

These highly consistent effects across tissue types remained when absolute volumes were decomposed into the individual structures within the Fischer 344 rat brain atlas. All 120 structures increased with linear age, 72 of 120 demonstrated a significant positive quadratic relationship with age (upward-facing curve), and 119 of 120 structures were larger in males. 85 of 120 structures demonstrated a significant negative linear age by sex interaction and 14 of 120 structures demonstrated significant positive quadratic age by sex interaction, increasing more with age in males than females. LME results and absolute volumes in mm^3^ for all structures can be found in **Supplementary Tables 3 and 4**, respectively.

### 3.2 Brain-size-corrected volumes show heterogeneous change with age and occur in regions implicated in motor control, learning, and memory

As shown in **Figure 1B** and **Supplementary Table 5,** aggregated brain-size-corrected volumes demonstrated different age-related effects depending on tissue type: total GM volume decreased significantly and demonstrated a positive curvilinear relationship with age (poly(age,2)1 β=-6.807, q=2.90E-17; poly(age,2)2 β=2.238, q=8.30E-04), while total WM volume increased sharply with linear age (poly(age,2)1 β=6.403 q=8.38E-09), and CSF volume increased subtly with linear age (poly(age,2)1 β=3.432, q=3.62E-02).

Aggregated volumes were then decomposed into the individual structures delineated by the Fischer 344 atlas. A visual representation of whole-brain age-related changes was performed using the *mincPlotSliceSeries* function in RMINC, wherein the t-statistics of LME modeling results split by model term were overlaid on the average anatomy background (**Figure 2**). LME results for selected structures, chosen for the strength of their effects or relevance of the brain region in aging and neurodegeneration (basal forebrain, caudoputamen, cingulum, commissure of the inferior colliculus, frontal cortex, dentate gyrus, hippocampus, internal capsule, nucleus accumbens, periaqueductal grey, and thalamus), are shown in **Table 1**. Visualization of the longitudinal trajectories for these selected structures, split by sex, was performed using the effects package in R (version 4.2-0, (Fox and Weisberg 2019)) and is shown in **Figure 3**. For simplicity and since effects were highly consistent across hemispheres, for structures that exist within both hemispheres, such as the hippocampus, only the trajectory within the right hemisphere is shown. LME results and volumetric trajectory visualization for all 120 structures are included in **Supplementary Table 5** and **Supplementary Figures 1 to 8**, respectively.

**Figure 2:**
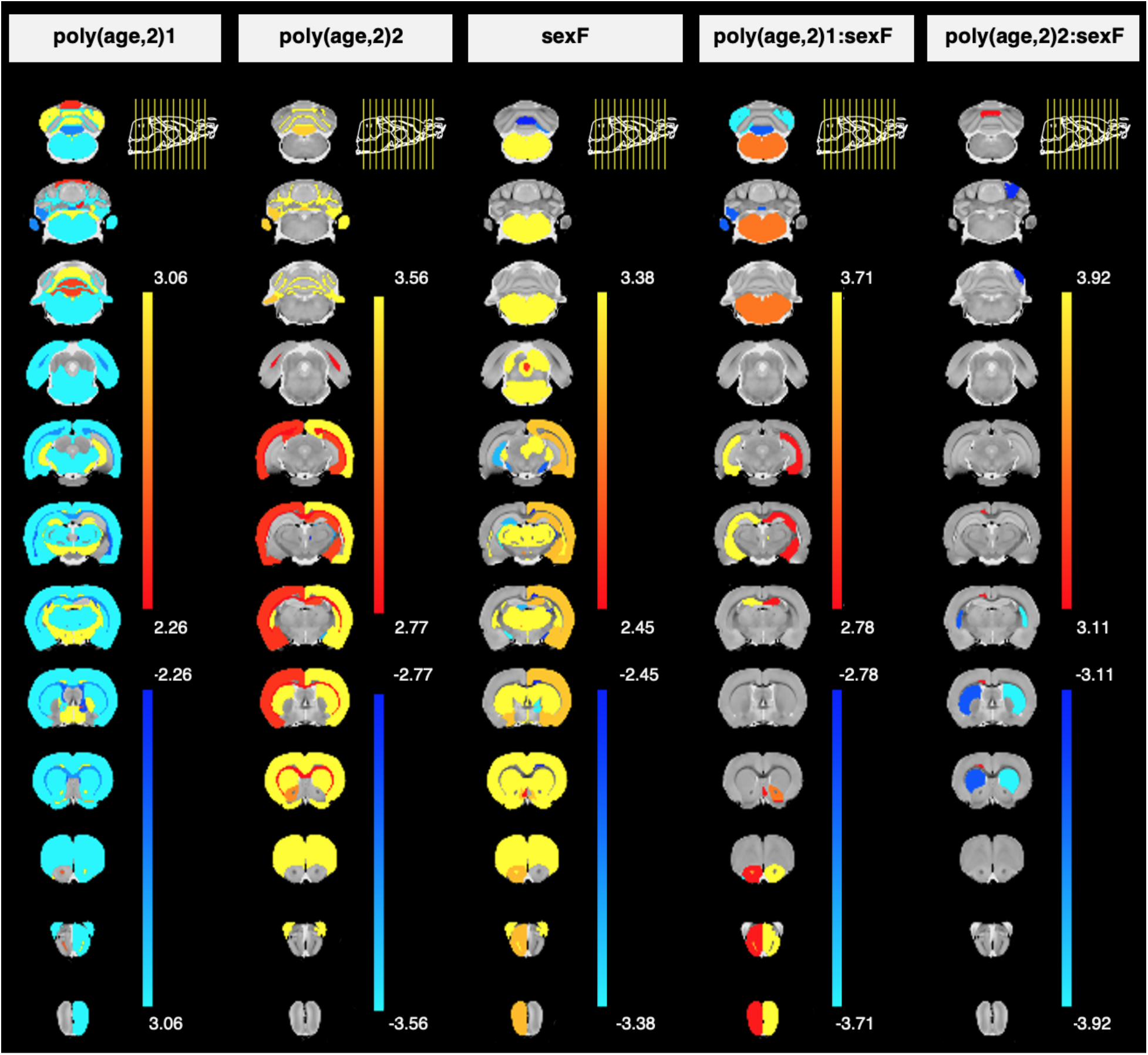
Visualization of age and sex effects, along with their interaction, on brain size-corrected brain volumes (LME results) in healthy aging Fischer rats, overlaid on the average anatomy background. In the mixed effects model, volumes were each predicted by an age by sex interaction with a random intercept for each subject. Age was modelled using a quadratic polynomial function to account for the possibility of non-linear changes with age. The linear and quadratic components of age are written as poly(age,2)1 or poly(age,2)2, respectively. Effects of sex (sexF) were evaluated in females relative to males, and the interaction between age and sex was also examined (poly(age,2)1:sexF and poly(age,2)2:sexF, with males as the reference group. The regional t-values for each term in the mixed effects model are indicated by the colour bars in columns 1 through 5. The plot range for each set of t-values is specific to each model term and displays effects significant between 5 and 1% FDR, with t-values above 1% thresholded at the 1% value, except for the poly(age,2)2:sexF column which displays values significant at 10% FDR.

**Figure 3:**
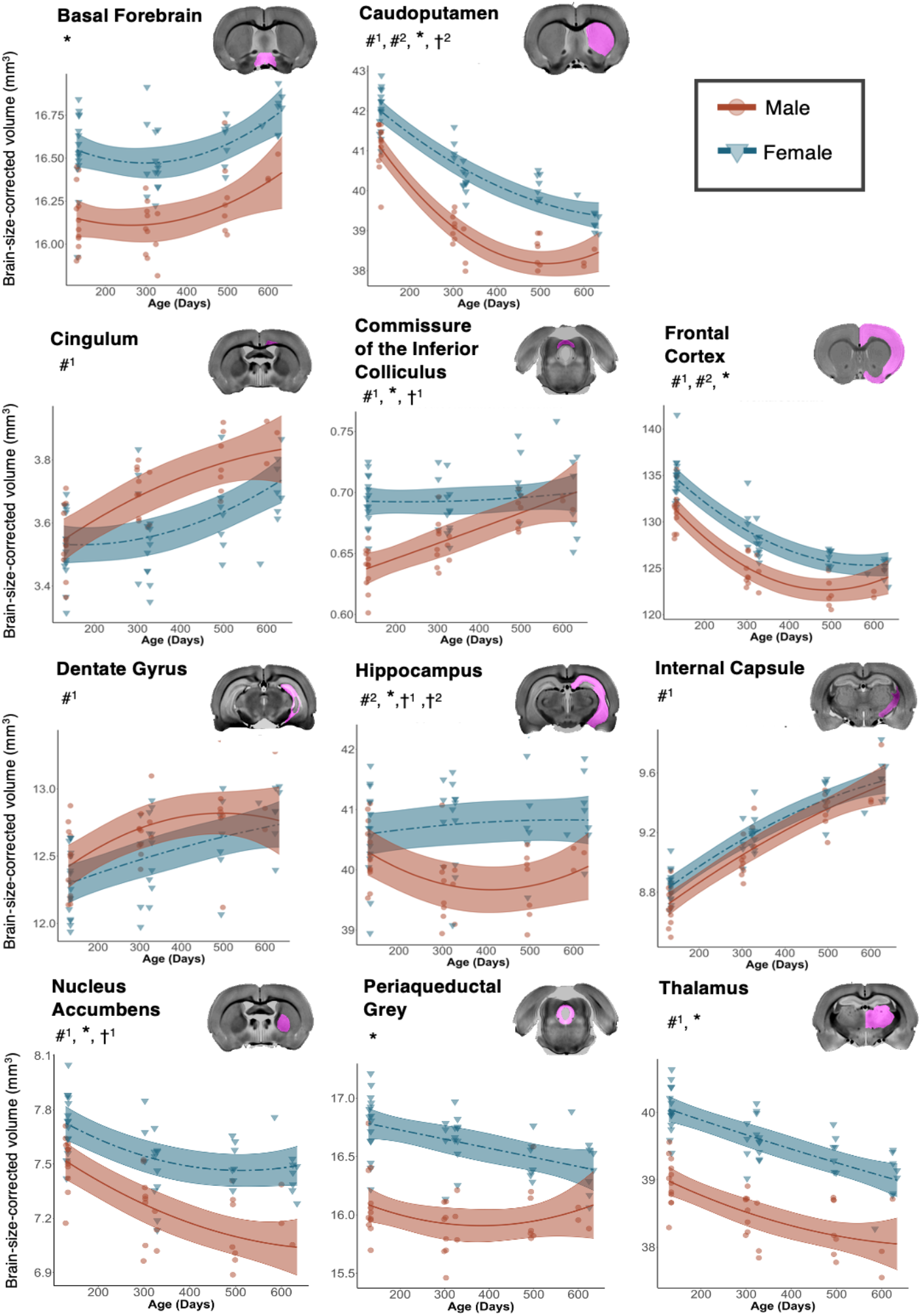
Prominent age and sex-dependent changes in brain-size-corrected volumes are present throughout the brain in both grey and white matter regions, suggesting a multitude of physiological functions are affected by aging. MRI data were acquired longitudinally in rats aged 4-, 10-, 16- and 20-months old. Main effects of age and sex, as well as age by sex interactions, were determined using linear mixed effects modelling. Each data point represents a single rat. The mixed effects model used to fit the data is represented by a line of best fit and 95% interval (shaded), split by sex. Data corresponding to males is shown using red circles with a solid line of best fit, while females are shown using blue triangles and a dashed line of best fit. Significance symbols are shown for each term in the model, where #^1^ and #^2^ represent the linear and second order age terms, * denotes a main effect of sex, and †^1^ and †^2^ represent age by sex interactions with a linear and second order age term, respectively. Multiple comparisons were corrected for using a 5% false discovery rate.

**Table 1:**
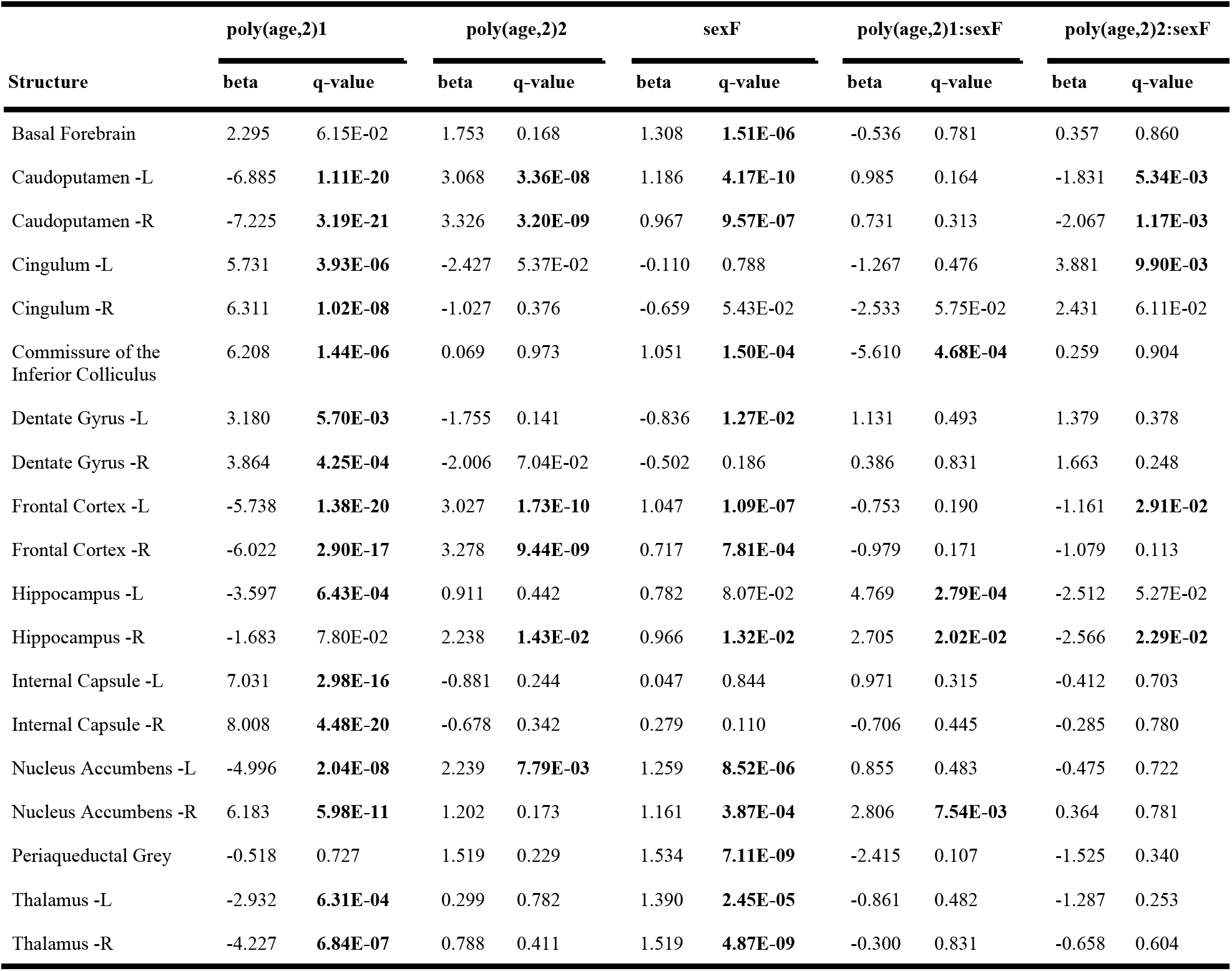
Linear mixed effects model summary for selected brain-size-corrected volumes, listed in alphabetical order. Volumes were predicted by an age by sex interaction with a random intercept for each subject. Age was modelled using a quadratic polynomial function. The linear and quadratic components of the age term with age are written as poly(age,2)1 and poly(age,2)2, respectively. Effects of sex were evaluated in females relative to males. Betas are standardized. Bold font denotes significance after FDR-correction at 5%. L or R represent the structure in the left or right hemisphere, respectively.

GM structures showed an almost even split of increased versus decreased volumes with age, while WM and ventricular system volumes generally increased with age. Specifically, of the 76 GM volumes analyzed, 20 regions increased linearly with age, while 24 decreased. 15 of 76 GM structures demonstrated a significant positive curvilinear relationship with age while 4 structures demonstrated a negative curvilinear relationship. Of the 40 WM volumes analyzed, 20 increased and 3 decreased linearly with age, while 4 of 40 regions demonstrated a negative curvilinear relationship with age. Of the 4 volumes that were categorized as part of the ventricular system, only the aqueduct and fourth ventricle increased significantly with age. Additionally, GM structures comprised the vast majority of the strongest age-related changes overall. Upon ranking the structures within the model terms poly(age,2)1 and poly(age,2)2 by q-value, of the top 20 strongest linear changes with age, 14 were in GM structures. For curvilinear changes the top 20 comprised 15 GM, 4 WM, and 1 CSF region(s). All structures ranked by q-value within each model term is shown in **Supplementary Table 6.**

The strongest linear changes with age in GM structures were in the temporal-parietal cortex (Left β=-6.904, q=4.29E-16; Right β=-6.937, q=1.02E-22), caudoputamen (Left β=-6.885, q=1.11E-20; Right β=-7.255 q=3.19E-21), hindbrain (β=7.611, q=3.19E-21), frontal cortex (Left β=-5.738, q=1.38E-20; Right β=-6.022, q=2.90E-17), and substantia nigra (Left β=7.113, q=8.70E-12; Right β=6.650, q=5.51E-12). Significant, but less strong age-related changes were also present in regions including the occipital-entorhinal cortex, nucleus accumbens, flocculus, thalamus, which all decreased, and the pons, hypothalamus, dentate gyrus, and crus 2 ansiform lobule, which all increased with linear age. In WM, the strongest linear changes with age were identified in the internal capsule (Left β=7.031, q=2.98E-16; Right β=8.008, q=4.48E-20), cerebral peduncle (Left β=6.857, q=4.87E-09; Right β=7.332, q=2.96E-14), and fimbria (Left β=5.575, q=3.41E-09; Right β=7.360, q=5.67E-12). Significant but less strong changes occurred in the cingulum, optic tract, stria terminalis, and commissure of the inferior colliculus, all of which increased with age. Age-related changes in CSF volumes were limited to the aqueduct (β=3.675, q=5.55E-03) and fourth ventricle (β=3.040, q=4.87E-02).

Regarding curvilinearity, the strongest effects in GM structures were in the frontal cortex (Left β=3.027, q=1.73E-10; Right β=3.278 q=9.44E-09), caudoputamen (Left β=3.068, q=3.36E-08; Right β=3.326, q=3.20E-09), and hindbrain (β=-3.309, q=6.19E-08). Curvilinear effects in WM structures were less common and were only significant in the optic chiasm (β=-3.419, q=5.24E-04), right optic tract (β=-2.007, q=5.88E-03), right fasciculus retroflexus (β=-3.261, q=2.08E-02), and left olfactory tract (β=-2.833, q=4.02E-02). Significant curvilinearity with age was not present in ventricular system volumes.

### 3.3 The influence of sex on neuroanatomy is strongest in grey matter structures

Of the brain-size-corrected volumes aggregated across tissue type, only total GM demonstrated significant sex-related effects. Total GM volume was significantly larger in females (sexF β=0.975, q=4.37E-09), as shown in **Figure 1B** and **Supplementary Table 5.** None of total GM, WM, or CSF volumes demonstrated age by sex interactions.

Upon examination of 120 regional volumes (split across hemispheres), GM structures comprised the majority of the sex effects identified in aging Fischer rats. 31 of 76 GM volumes showed significant effects of sex, with 23 of 31 being larger in females. 13 of 40 WM volumes had significant sex effects, with 9 of 13 being larger in females. Additionally, Of the top 20 smallest (most significant) q-values, 13 were in GM structures and 7 in WM structures (**Supplementary Table 6**). The strongest main effects of sex in GM structures were in the caudoputamen (Left β=1.186, q=4.17E-10; Right β=0.967, q=9.57E-07), thalamus (Left β=1.390 q=2.45E-05; Right β=1.519, q=4.87E-09), periaqueductal grey (β=1.534, q=7.11E-09), and basal forebrain (β=1.308, q=1.51E-06), with weaker changes occurring in the nucleus accumbens, globus pallidus, and frontal cortex, all of which were larger in females. The most significant main effects of sex in WM structures were in the optic chiasm (β=-1.125, q=3.85E-05) and commissure of the inferior colliculus (β=1.051, q=1.50E-04), with additional effects in the stria medullaris of the thalamus, mammillothalamic tract, and trochlear nerve, wherein all were larger in females relative to males.

Finally, age by sex effects were present in grey and white matter volumes, but not the ventricular system; 14 of 76 GM structures and 8 of 40 WM structures demonstrated linear age by sex interactions, with a positive interaction occurring in 8 of 14 grey matter regions, and 6 of 8 white matter regions, reflecting increasing volume in females relative to males as a function of age. A second order age by sex interaction was present for 8 of 76 grey matter regions and 3 of 40 white matter regions; this interaction was negative for 7 of 8 grey matter regions, reflecting a more negative curvilinear relationship with age in females than in males, while all 3 white matter regions showed the opposite effect, demonstrating positive curvilinear relationships with age.

Significant age by sex interactions were generally observed bilaterally, though with differing strengths of effect. The linear interaction between age and sex in GM structures was strongest in the crus 2 ansiform lobule (Left β=-5.173, q=1.62E-04; Right β=-6.824, q=1.79E-07), hippocampus (Left β=4.769, q=2.79E-04; Right β=2.705, q=2.02E-02), olfactory nuclei (Left β=3.404, q=1.94E-02; Right β=5.150, q=1.14E-03), and cerebellar lobule 7 (β=-5.044, q=1.19E-03). Additional significant effects were identified in the left copula and cerebellar lobule 8, which demonstrated negative age by sex interactions, and the right nucleus accumbens and right ventral pallidum, which showed positive age by sex interactions. In WM structures a linear age by sex interaction was present in fewer structures and included the left lateral olfactory tract (β=-6.786, q=6.90E-05), commissure of the inferior colliculus (β=-5.610, q=4.68E-04), optic chiasm (β=3.952, q=1.28E-03), intrabulbar part of the anterior commissure (Left β=3.719, q=5.31E-03; Right β=3.200, q=2.49E-02), and optic tract (Left β=2.120, q=4.31E-02; Right β=2.020, q=3.07E-02).

Finally, second order age by sex interactions in GM structures were strongest in the caudoputamen (Left β=-2.566, q=2.29E-02; Right β=-1.831 q=5.34E-03), right hippocampus (β=-2.566, q=2.29E-02; the left hippocampus neared significance with β=-2.512, q=5.27E-02), and left frontal cortex (β=-1.161, q=2.91E-02), with additional effects in the right simple lobule, crus 1 ansiform lobule, left occipital-entorhinal cortex, and cerebellar lobule 4/5, wherein all but the occipital-entorhinal cortex demonstrated negative curvilinear relationships with age. In WM structures the only significant quadratic age by sex interactions were in the left cingulum (β=3.881, q=9.90E-03), optic chiasm (β=3.007, q=1.32E-02), and right lateral olfactory tract (β=3.785, q=3.84E-02).

### 3.4 Whole-brain voxelwise and volumetric analyses provide complementary results

A whole-brain voxelwise analysis was performed in conjunction with the volumetric analysis to explore if the two methods would provide complementary information. A visual representation of whole-brain age- and sex-related voxelwise changes was performed using the *mincPlotSliceSeries* function in RMINC, wherein the t-statistics of LME modeling results at each voxel were overlaid on the average anatomy background, split by model term (**Figure 4**). Side-by-side comparisons of volumetric and voxelwise results for the effects of age and the influence of sex (main effect and interaction with age) are shown in **Supplementary Figures 9 and 10,** respectively. Overall, voxelwise results correspond well with those identified using volumetric analyses. However, the volumetric analysis—whereby the relative Jacobians shown in the voxelwise data were integrated across each structure in the Fischer atlas—was able to identify changes with age or between sexes that were not visible at the individual voxel level, and similarly, several focal changes exist that were not visible at the whole-structure level.

**Figure 4:**
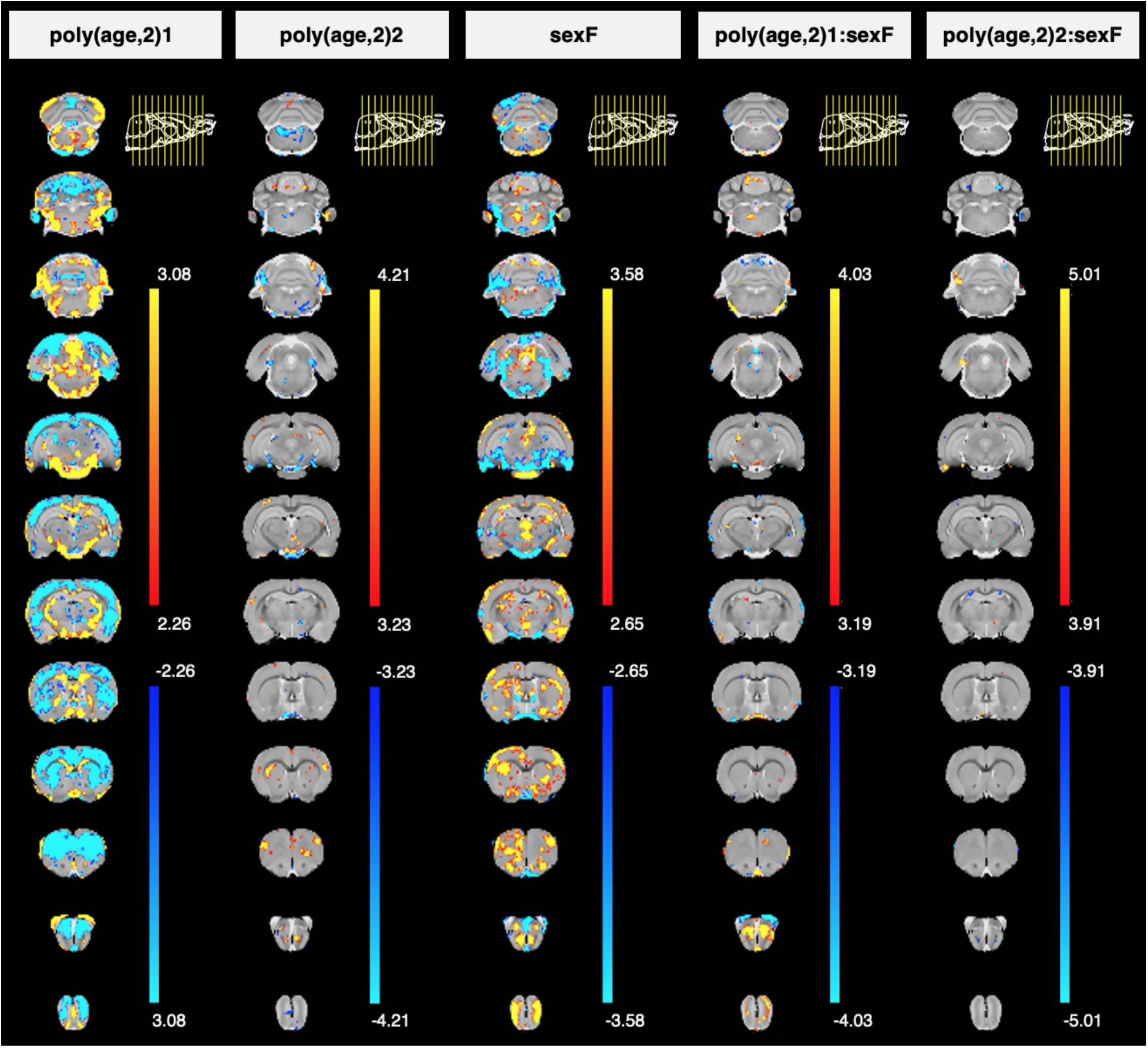
Statistical maps demonstrating local brain size-corrected volume differences due to age, sex, and their interaction in healthy aging Fischer rats, overlaid on the average anatomy background. A mixed effects model was run at each voxel across the brain, whereby relative Jacobians were predicted by an age by sex interaction with a random intercept for each subject. Age was modelled using a quadratic polynomial function to account for the possibility of non-linear changes with age. The linear and quadratic components of age are written as poly(age,2)1 or poly(age,2)2, respectively. Effects of sex (sexF) were evaluated in females relative to males, and the interaction between age and sex was also examined (poly(age,2)1:sexF and poly(age,2)2:sexF, with males as the reference group. The t-values for each term in the mixed effects model are indicated by the colour bars in columns 1 through 5. The plot range for each set of t-values is specific to each model term and displays effects significant between 5 and 1% FDR, with t-values above 1% thresholded at the 1% value.

The linear effect of age was generally consistent between voxelwise and volumetric analyses, though with a few notable differences. In the cerebellum, the voxelwise analysis identified both significant focal increases and decreases within cerebellar lobule 4 and 5. These changes were offset when summing the volume over the whole region, resulting in a lack of linear age effect in the volumetric analysis. Further, the voxelwise analysis revealed consistent significant increases in the medial part of the corpus callosum, while much of the lateral corpus callosum saw significant decreases, resulting in a net decrease in relative corpus callosum volume when analyzing at a regional level.

When examining the second order effects of age, the most noticeable contrast between the two methods was throughout the cortex. While the voxelwise analysis failed to identify second order effects of age, the volumetric analysis identified highly significant positive curvilinear effects in cortical volumes with age.

Regarding the main effect of sex, there were several brain regions whereby the voxelwise analysis identified focal increases and decreases in females relative to males within the same structure. These contrasting focal effects appeared to neutralize each other within the regional analysis, resulting in a lack of overall effect. This is particularly clear in the cerebellar lobules, whereby the voxelwise analysis identifies significant sex-related increases in the posterior cerebellar lobule 3, and increases in the anterior cerebellar lobule 3, resulting in a lack of overall effect at the level of the whole structure.

When exploring age by sex interactions there was further evidence of complementary information gained by using both voxel-based and volumetric analyses. Regarding linear age by sex interactions, there were positive bilateral effects in the hippocampal formations that were not strongly reflected at a voxelwise level. Focal effects also drove whole-volume effects; the strong positive interaction of linear age and sex in hindbrain volume appeared to be driven by focal changes within the medial hindbrain, though the specific nuclei were not identifiable given resolution and contrast of the images. Finally, upon comparing the effects of the interaction of curvilinear age and sex identified by each method, there were bilateral decreases in the caudoputamen at a volumetric level, which were not reflected at a voxelwise level.

## 4. DISCUSSION

The purpose of this study was to characterize brain volumetric changes and whole-brain voxelwise changes in a commonly used rat model of aging (Gallagher, Stocker, and Koh 2011). This work establishes a baseline for normative neuroanatomical change with age in the rat brain, and informs studies examining pathology-related change in transgenic models developed on the Fischer 344 background strain. It is particularly important to study both sexes in this context, given the progression and presentation of many age-related diseases, such as Alzheimer’s disease, are influenced by sex. Our study is among the few aging studies to assess structural change longitudinally, allowing for exploration of non-linear effects and providing increased power to detect small differences in neuroanatomy.

119 of 120 absolute volumes increased linearly with age, with 72 of 120 structures also demonstrating a positive curvilinear relationship with age. These trends were also seen in the total aggregated volumes for GM, WM, and CSF. This consistent volumetric increase with age matches the few studies that have tracked absolute change in regional and/or total brain volume with age in wildtype rodents (Casas et al. 2018; Gaser et al. 2012; Maheswaran et al. 2009). Our finding that 119 of 120 brain structures were larger in males when measured using absolute volume can be explained by the documented relationship between brain size and body size (Welniak-Kaminska et al. 2019; Valdés-Hernández et al. 2011), and matches previous studies comparing absolute brain volumes between males and females quantified at various periods during the rodent lifespan (Sumiyoshi, Nonaka, and Kawashima 2017; Qiu et al. 2018; Spring, Lerch, and Henkelman 2007; Reichel et al. 2017), and in humans (Scahill et al. 2003; Walhovd et al. 2005; Jay N. Giedd et al. 2012). Interestingly, this significant progressive increase in brain volume throughout the Fischer rat lifespan is in contrast to that in humans, which plateaus in the mid-teens and then declines later in life (J. N. Giedd et al. 1999). The difference between rodent and human brain growth is generally hypothesized to be facilitated by the delay in growth plate closure in rodents, permitting continuous expansion of brain volume (Sandner et al. 2010; Kilborn, Trudel, and Uhthoff 2002).

Because of inherent differences in total brain size between sexes and the fact that absolute volume in rodents increases across the majority of brain regions with age, quantification of absolute volumes is not particularly informative. However, the absolute volume in mm^3^ of 73 brain regions, split by timepoint, hemisphere, and sex (**Supplementary Table 4**), may still be useful to the preclinical neuroimaging community since it can be used as *a priori* information for power analysis for future MRI-based studies in adult rats.

The analysis of brain-size-corrected volumes offers a much more interesting and relevant perspective on how the brain is altered locally during normal aging and between sexes. In the majority of other studies examining volumetric change this is done by normalizing (dividing) regional volumes to intracranial volume (ICV) or total brain volume (TBV). We employed an alternative method to explore volumetric change that avoids the inherent alteration of the distribution of variance, as well as the increase in measurement uncertainty that occurs when error propagation is performed, as is necessary when dividing a regional volume by ICV or TBV. Our corrected volumes were obtained by integrating relative Jacobians across voxels within a given structure, thus providing a value representative of local regional change independent of affine scaling factors for global brain volume, while avoiding violating assumptions regarding the distribution and variance incurred by computing a ratio of total volume.

The age-related changes in total GM, WM, and CSF volumes in the Fischer rat brain that we report are generally consistent with the few existing aging studies in rodents that examine relative volumetric change. Given the overall lack of rodent aging studies examining volumetric change and the desire to characterize the translatable nature of this work as it pertains to humans, we also briefly compare our findings to those reported in human aging studies.

Total CSF compartmental volume increased with age, as has previously been documented in both rodents (Maheswaran et al. 2009; von Kienlin et al. 2005) and humans, particularly in the lateral ventricles (Fjell et al. 2009; Scahill et al. 2003; Driscoll et al. 2009; Walhovd et al. 2005; Narvacan et al. 2017). Regionally, only the aqueduct and fourth ventricle increased significantly with age, indicating the majority of ventricular enlargement occurred in the more caudal parts of the Fischer 344 rat brain.

Total WM volume increased linearly in aging Fischer 344 rats. When separated into individual WM structures, linear changes with age were identified, with few curvilinear effects. The strongest linear changes with age were identified in the internal capsule, cerebral peduncle, and fimbria, with significant but less strong changes occurred in the cingulum, optic tract, stria terminalis, and commissure of the inferior colliculus, all of which increased. These structures are spatially distributed across the brain and are implicated in a number of physiological functions, including motor function and integration of visual-motor information ((internal capsule, (Wen et al. 2019), cerebral peduncles, (E. W. Jenkinson and Glickstein 2000; Howzet 1974)), integration of bilateral information in the auditory system (commissure of the inferior colliculus, (Lee, Yanagawa, and Imaizumi 2015)), episodic and spatial memory formation (fimbria, cingulum, (Bubb, Metzler-Baddeley, and Aggleton 2018; Aggleton and Brown 1999; Okada and Okaichi 2006)), integration of emotional information such as fear, anxiety, and stress (stria terminalis, (Lebow and Chen 2016)), and transmission of visual information (optic tract, (Mehra and Moshirfar 2020; Calkins 2013)).

In humans, WM volume appears to increase until approximately age 45 and then plateaus and decreases (Hedman et al. 2012; Hasan et al. 2010; Walhovd et al. 2005; Narvacan et al. 2017; Pfefferbaum et al. 2013), in opposition to our finding that total WM increased consistently with age. Unfortunately our ability to understand this discrepancy is limited by the lack of volumetric analyses of the majority of white matter regions, particularly at the preclinical level. As such, it is difficult to know if the continued age-related increase in these regions and in total WM after mid-life is unique to Fischer 344 rats, or to rodents in general. One longitudinal study in C57BL6 mice demonstrated significant age-related volumetric increases in the corpus callosum, corticospinal tract, and the fornix system up until 14 months of age in WT mice, but did not perform scans later into the lifespan (Maheswaran et al. 2009). An explanation for the consistent age-related increase in white matter volume could be that Fischer rats display WM decline much closer to the end stages of their lifespan; Fischer rats live 21 months on average but they can live up to 26 to 30 months, though with an attrition rate of 75 to 94%, respectively (Chesky and Rockstein 1976). An alternate interpretation is simply that the strong age-related increase in white matter volume seen in our aging Fischer rats (and C57BL6 mice, (Maheswaran et al. 2009) represents a fundamental difference between human and rodent brains when it comes to hallmarks of normative aging. Additional longitudinal studies in rodents that extend late into their lifespan and that document both whole-brain and regional tissue-specific change are clearly needed to characterize age-related change in WM volumes, and to better understand differences in WM changes between aging rodents and humans.

Total GM volume decreased in aging Fischer 344 rats, and also demonstrated a positive curvilinear relationship with age, similar to the linear and curvilinear decreases in GM volume in humans post-adolescence that have been consistently shown in human aging studies (Hedman et al. 2012; Hasan et al. 2010; Walhovd et al. 2005; Narvacan et al. 2017; Pfefferbaum et al. 2013).

Our age-related findings within specific GM structures closely match those reported by other authors at the preclinical and clinical level. The most relevant example is a longitudinal study in male C57BL6 mice from 6 to 14 months of age that reported slight, non-significant decreases in the cortex, hippocampus, caudoputamen, and thalamus, and increases in the midbrain-hindbrain and hypothalamus ((Maheswaran et al. 2009). We also report age-related effects in these regions, though the decreases that we identified are highly significant, possibly because our study explored changes 6 months later into the lifespan. The increases they identified in the midbrain-hindbrain and hypothalamus of aging C57BL6 mice are similar to those seen in our Fischer rats.

Regarding other regional changes within the brain, the only other study to examine relative change in wildtype rodents comparably late in life is a cross-sectional voxel-based morphometry study in Fischer rats at 10 and 25 months (Alexander et al. 2020). Interestingly, the authors report a relative increase in select hippocampal regions, which, upon comparison to the rat atlas used in the present study, appears to be primarily within the dentate gyrus, a region in which we identified increased volume with age in both our volumetric and voxel-based analyses. In contrast, decreased dentate gyrus volume in the rat brain has previously been reported (A. K. Shetty and Turner 1999), which matches the majority of human aging literature (Small et al. 2004; Dillon et al. 2017; Hayek et al. 2020; Wisse et al. 2014; Malykhin et al. 2017). Additionally, some of the changes that we document here, such as decreased volume in the substantia nigra, nucleus accumbens, mammillary bodies, flocculus, and paraflocculus, and increased pons and crus II ansiform volume, have yet to be reported in preclinical literature. Given the sparsity of literature on relative volumetric change with age in the rodent brain, additional studies are required to improve the interpretation of our findings.

Although several of our findings have not yet been reported in preclinical literature, there are some similarities compared to human aging literature. Aging studies in humans have identified highly heterogeneous age-related change across the brain, including atrophy in many subcortical gray matter structures, including the thalamus, nucleus accumbens, caudate, and putamen, (Long et al. 2012; Walhovd et al. 2005; Narvacan et al. 2017; Tullo et al. 2019), and decreased cerebellar volume (Bernard and Seidler 2013; Han et al. 2020), all of which we also report in our aging Fischer rats. Hypertrophy of the substantia nigra (Cabello et al. 2002; Rudow et al. 2008), and either stable or increased hippocampal volume until mid-life followed by a sharp decline have also been reported (Narvacan et al. 2017; Malykhin et al. 2017; Raz et al. 2010; Bussy et al. 2021), again, similar to our findings. For reviews on age-related structural changes in humans see (Sowell, Thompson, and Toga 2004) and (Fjell and Walhovd 2010).

Regarding the cellular and molecular basis for altered brain structure with age, it has previously been shown that altered axonal/dendritic branching, synapse, spine, or cell numbers are sufficient to alter brain volume (J. P. Lerch et al. 2011; Qiu et al. 2013; Spring et al. 2010). Our study design did not allow us to identify these underlying mechanisms, so we refer readers to publications that have explored these topics in detail (Morterá and Herculano-Houzel 2012; Rapp and Gallagher 1996; Uylings and de Brabander 2002; Mattson and Arumugam 2018; Dumitriu et al. 2010). Additionally, we recently published a longitudinal analysis of hippocampal neurochemical concentrations in the same cohort of aging rats as studied here (Fowler et al. 2020). We identified altered concentrations of metabolites implicated in neuroinflammation, bioenergetics, and antioxidant capacity, which, while localized to the hippocampus, are indicative of major changes at the cellular level during normal aging.

Structures influenced by sex were widespread across the Fischer rat brain and more commonly found in GM than WM structures, for both the main effect of sex and its interaction with age. Previous explorations of the influence of sex on brain-size-corrected or relative volumes have been performed at the preclinical level, though with limited comparability to our findings. In preclinical studies, sex differences in relative brain volumes have not been examined in the context of aging, but rather, during brain development (6-10 weeks, (Sumiyoshi, Nonaka, and Kawashima 2017); post-natal day 3-65, (Qiu et al. 2018)), or at a single time point in young adulthood (12 weeks, (Spring, Lerch, and Henkelman 2007)).

In our cohort of aging Fischer rats, females displayed larger volumes in the frontal cortex, caudoputamen, thalamus, and periaqueductal grey relative to males, similar to findings by Qui et al. in C57BL/6J mice, while in contrast, Spring et al. noted smaller thalamic volumes in adolescent female Wistar rats. The influence of sex on hippocampal volumes in rodents is also inconsistent. We identified a positive linear age by sex interaction in hippocampal volume, indicating larger hippocampi in female rats relative to males across the life span, while the opposite has been reported in Wistar rats (Sumiyoshi, Nonaka, and Kawashima 2017). Spring et al. reported a larger anterior hippocampus but smaller posterior hippocampus in female C57BL/6J mice (Spring, Lerch, and Henkelman 2007). Cerebellar volumes have been reported to be larger in female mice (Qiu et al. 2018), though smaller in female rats relative to males (Sumiyoshi, Nonaka, and Kawashima 2017), which matches our findings.

Given the lack of studies examining the influence of sex in relative brain volumes in rodents at ages comparable to those studied here, it is difficult to determine if discrepancies between our findings and the three aforementioned studies are due to inherent differences between mouse and rat brains, differences between strains of rats, or in part due to the younger age of the rodents studied by the other authors compared to our cohort of Fischer rats. As such, many of our findings require additional preclinical research for confirmation, particularly the sex effects within the basal forebrain, crus 2 ansiform lobule, cingulum, commissure of the inferior colliculus, ventral pallidum, intrabulbar part of the anterior commissure, and optic chiasm, since this is the first time, to our knowledge, that these effects have been documented in the aging rodent brain. In order to properly detect the influence of sex on brain structure in future studies, it will be important to ensure the rodent brain template used is equally representative for males and females, as is the case with the Fischer rat atlas we developed in our lab and employed here (Goerzen et al. 2020).

Comparisons to human literature examining the influence of sex on neuroanatomy further confirm several of our findings. In healthy adults across the lifespan, total grey matter has been shown to be larger in females after controlling for intracranial volume (Cosgrove, Mazure, and Staley 2007; Armstrong et al. 2019; Leonard et al. 2008), as has overall cortical volume (Goldstein et al. 2001), consistent with our study. Sex differences in rates of normal age-related change have been observed for ventricular CSF and cortical thickness and/or volume within the middle frontal, parahippocampal, and superior parietal regions, whereby change with age was more pronounced in males (Driscoll et al. 2009; Cowell et al. 1994; Coffey et al. 1998; Armstrong et al. 2019; Pfefferbaum et al. 2013; Sullivan et al. 2004). In direct contrast, we report similar age-related change in cortical and ventricular volumes in male and female Fischer rats, though with some additional curvilinearity with age in the male trajectories.

There are only a few subcortical structures for which sex differences during aging have been identified, though plenty of studies exist on sex differences in brain structure outside of the context of aging. For reviews see (Cosgrove, Mazure, and Staley 2007; Jay N. Giedd et al. 2012; Ruigrok et al. 2014). It has previously been reported that women have larger hippocampal volumes (Cosgrove, Mazure, and Staley 2007; Jay N. Giedd et al. 2012; Goldstein et al. 2001), while men have been shown to have steeper age-related volumetric declines (Armstrong et al. 2019), similar to our identification of larger hippocampal volume in females with atrophy occurring primarily in male rats. We also report larger thalamic volume in female Fischer rats, with atrophy occurring at a similar rate in males and females, comparable to human aging cohorts (Ruigrok et al. 2014; Sullivan et al. 2004). Caudate and/or putamen volume has been shown to be larger in women (Cosgrove, Mazure, and Staley 2007; Jay N. Giedd et al. 2012), similar to our finding of larger caudoputamen volume in female Fischer rats, but to decline similarly with age in both males and females (Choi et al. 2020; Murphy et al. 1996), while we report a steeper decline and then slight recovery in volume in aging males. Overall, sex-related structural changes in aging Fischer rats generally align with what has been reported in the human literature, though many of our findings could not be compared, particularly those in WM structures, due to a lack of literature exploring the influence of sex on many of the structures we studied.

The origin and subsequent impact of sex chromosomes and hormones on the brain have been studied at the cellular, molecular, structural, and behavioural level. For reviews, see (Cooke et al. 1998; Cahill 2006; Osterlund and Hurd 2001). It has been shown that areas of the brain with high levels of sex steroid receptors during periods of brain development show greater sexual dimorphisms, such as the precentral, superior frontal, and lingual gyri, hippocampus, caudate, and thalamus (Goldstein et al. 2001; Qiu et al. 2018). Sex hormones influence the outgrowth of axons and dendrites, the amount of cell death, and the number and type of synapses that a cell makes (Juraska, Sisk, and DonCarlos 2013; Juraska and Lowry 2012), all of which are sufficient to alter brain volume (J. P. Lerch et al. 2011; Qiu et al. 2013; Spring et al. 2010). Recent studies using Four Core Genotype mice (which can be gonadally male or female with XX or XY chromosomes), or X-monosomic mice (XO as opposed to XX) have even allowed researchers to decouple the impact of sex chromosomes from gonadal hormones on brain structure, as well as spatial learning and memory (Corre et al. 2016; Raznahan et al. 2013).

Regarding the influence of sex on brain structure during aging, previous studies have shown that in addition to hormone levels fluctuating with age, topographic distribution, binding capacity, and associated enzyme levels of sex steroid receptor systems also vary as a function of age (Sholl and Kim 1990; MacLusky et al. 1987), likely contributing to differing trajectories of aging in males and females in specific regions. Specifically, there is evidence that brain regions with high androgen (and low estrogen) receptor populations early in development may be more susceptible to aging effects (Sholl and Kim 1990; Cowell et al. 1994) and that female sex hormones may protect the brain from age-related atropy (Gur et al. 1991). However, the role of sex hormones and their receptor systems in the living human brain have not been yet studied. Until this kind of data is collected, ensuring proper age and sex-matching in study design will be important, as the research community works towards understanding the influences of sex on age- and pathology-related changes in brain structure.

Finally, a whole-brain voxelwise analysis was performed in conjunction with the volumetric analysis to explore if the two methods would provide complementary information. Overall, the voxelwise and regional analyses reveal comparable findings. However, there are instances where the regional analysis masked some finer-grained details of morphological change identified in the voxelwise analysis, while conversely, analysis at the level of whole structures was capable of identifying changes that were only weakly present at the individual voxel level. This was particularly noticeable when exploring curvilinear effects of age, whereby quadratic age effects in the cortex were absent at the voxel level, but strongly present within the regional analysis. Conversely, contrasting focal sex effects were present in the cerebellar lobules, which negated each other in the regional analysis. As such, the inclusion of two methods for morphometric analysis enabled identification of changes that were complementary between the two methods.

This study has a few important limitations that warrant discussion. First, due to some of the rats being part of a treatment study at 10 months and therefore not included in this analysis, the number of animals decreased considerably at the last two time points. This may have decreased our power to detect differences between sexes or change with age. To confirm the robustness of the findings presented here, we performed the same analysis for the brain-size-corrected volumes across only three time points as opposed to four, compared side by side in **Supplementary Table 7**. The majority of age and sex effects remained in the three time point analysis, particularly the main effects of sex. Specifically, of the 179 significant findings in the original analysis, 132 were maintained, with effects lost in 23 regions (of these, 7 remained sub-threshold of significance, with q-values between 0.05 and 0.1), and significance gained in 24 regions (12 were previously subthreshold), primarily due to changes in curvilinearity. Despite changes in significance, the direction of change (sign of beta coefficient) was unaltered for all effects. Overall, while it is clear that the last time point at 20 months is important for consolidating curvilinear change, the majority of the results are consistent across the two analyses, demonstrating the robustness of our findings despite the decreased sample size towards the end of the study. The longitudinal nature of the study, the number of time points over which data were collected, and the implementation of a 5% false discovery rate correction also improve the confidence with which we report our findings.

Additionally, the image resolution in this study (114 μm) prevented the delineation of very small structures and/or adjacent grey matter nuclei, such as the thalamic, cortical, and amygdalar nuclei. To address this limitation, we performed a whole-brain voxel-wise analysis to identify focal changes that may not have been identified in the volumetric analysis, which confirmed that the two methods revealed complementary information regarding neuroanatomical change, and even found some focal effects masked by regional averaging.

## 5. CONCLUSION

In order to better understand and address age-related pathologies in transgenic models of disease, it is necessary to first characterize the features of normal aging in common experimental background strains, such as the Fischer 344 wildtype rat. To this end, this work presents a comprehensive analysis of MRI-derived brain changes at the voxelwise and whole-structure level during normative aging in a mixed-sex cohort of F344 rats. These findings contribute to our understanding of the neuroanatomical changes associated with normal aging in male and female F344 rats, critical for informing future studies in transgenic models of age-related diseases, which frequently present and progress differently in males versus females.

## Supporting information

Supplementary Images 1-11

## SUPPLEMENTARY MATERIAL AND METHODS

### Methods

#### Age- and sex-dependent change in brain-size-corrected volumes over three time points as compared to four time points

To ensure our results were not being driven by the last time point wherein we have the fewest subjects (and therefore, the least power), we also analyzed all data from the first three time points only. 68 of 70 regions that originally indicated linear effects of age maintained those effects over three time points as opposed to four, while five new regions demonstrated significant linear effects. Curvilinear effects were seen in 20 regions as opposed to 22 in the original analysis, losing significance in six regions but becoming significant in four. The main effects of sex remained particularly strong, with only three regions losing significance when analyzed over three timepoints. Four additional regions showed linear age by sex interactions, while one region was no longer significant. Quadratic age by sex interactions differed somewhat (nine new regions, seven regions no longer significant, four remained the same), which was to be expected given the aforementioned differences in curvilinear effects of age over three timepoints versus four. For a comparison of LME results obtained using data from three versus four time points see **Supplementary Table 6**. This additional analysis represents the steps taken towards ensuring our results are robust, despite the decreased sample size towards the end of the study. It is clear that the last timepoint at 20-months constitutes an important data pointsolidifying the curvilinear effects of age, and especially age by sex interactions, in brain structure present between 4 and 16 months.

### Tables and Figures

**Supplementary Table 1: Subject Data.** Number of subjects scanned at each timepoint, split by sex, with reasons for removal of scans indicated. -QC represents the number of scans removed due to failing quality control (QC), D represents death of a rat prior to that time point, and Tx denotes the removal of a rat after the 10-month time point due to participation in a separate treatment study. F and M denote female or male rats, respectively.

**Supplementary Table 2: Comparison between models containing a linear versus quadratic age term using Akaike information criterion (AIC).** In both models, volume was predicted using an age by sex interaction with a random effect per subject. In model 1, the age term was modelled using a linear age term (poly(age,1)) while in model 2, a quadratic age term (poly(age,2)) was used. Both used the polynomial function, but with differing degrees, to ensure proper model nesting for AIC comparison. Δ_i_ (AIC_i_-AIC_min_) was calculated using the AIC value for each model, whereby a Δ_i_ ≥ 4 indicated substantially less support for model 1 (first order age term) than model 2 (second order age term). The smallest AIC between model 1 and 2 is shown in grey, while Δ_i_ ≥ 4 and AIC_min_ are denoted by shaded blue boxes.

**Supplementary Table 3: Linear mixed effects model results for 73 absolute volumes derived from integrated absolute Jacobians.** Absolute volumes were modelled using an age by sex interaction with a random effect per subject. Age was expressed using a polynomial function of degree 2 to allow for non-linear change with age. The linear and quadratic components of the age term are written as poly(age,2)1 and poly(age,2)2, respectively. Effects of sex (sexF) were evaluated in females relative to males, along with the interaction of age and sex (poly(age,2)1:sexF and poly(age,2)2:sexF). Beta represents the standardized beta or coefficient value for each model term, with the adjacent column, Std.Error indicating the error associated with the standard beta measurement. The t-value for each model term is also included. Significant q-values, after 5% FDR correction, are denoted by blue shaded boxes, while those between 5 and 10% are shown by shaded grey boxes. L and R indicate a structure that is split over the left and right hemispheres, and was therefore reported and analyzed as two separate volumes.

**Supplementary Table 4: Absolute volumes of 73 brain regions in mm^3^, split by timepoint and sex.** Average volume and standard deviation for each structure across all subjects are recorded, with separate columns for the same data split by sex. L and R indicate a structure that is split over the left and right hemispheres, and was therefore analyzed and reported as two separate volumes.

**Supplementary Table 5: Linear mixed effects model results for 73 brain-size-corrected volumes derived from integrated relative Jacobians.** Brain-size-corrected volumes were modelled using an age by sex interaction with a random effect per subject. Age was expressed using a polynomial function of degree 2 to allow for non-linear change with age. The linear and quadratic components of the age term are written as poly(age,2)1 and poly(age,2)2, respectively. Effects of sex (sexF) were evaluated in females relative to males, along with the interaction of age and sex (poly(age,2)1:sexF and poly(age,2)2:sexF). Beta represents the standardized beta or coefficient value for each model term, with the adjacent column, Std.Error indicating the error associated with the standard beta measurement. The t-value for each model term is also included. Significant q-values, after 5% FDR correction, are denoted by blue shaded boxes, while those between 5 and 10% are shown by shaded grey boxes. L and R indicate a structure that is split over the left and right hemispheres, and was therefore reported and analyzed as two separate volumes.

**Supplementary Table 6: Linear mixed effects model results for 73 unique brain-size-corrected volumes derived from integrated relative Jacobians, ranked by q-value within each model term.** Brain-size-corrected volumes were modelled using an age by sex interaction with a random effect per subject. Age was expressed using a polynomial function of degree 2 to allow for non-linear change with age. The linear and quadratic components of the age term are written as poly(age,2)1 and poly(age,2)2, respectively. Effects of sex (sexF) were evaluated in females relative to males, along with the interaction of age and sex (poly(age,2)1:sexF and poly(age,2)2:sexF). Beta represents the standardized beta or coefficient value for each model term, with the adjacent column, Std.Error indicating the error associated with the standard beta measurement. The t-value for each model term is also included. Significant q-values, after 5% FDR correction, are denoted by blue shaded boxes, while those between 5 and 10% are shown by shaded grey boxes L and R indicate a structure that is split over the left and right hemispheres, and was therefore reported and analyzed as two separate volumes. Structures are ranked by q-value, from smallest (most significant) to largest.

**Supplementary Table 7: Linear mixed effects model results for brain-size-corrected volumes analyzed over 3 time points as compared to four.** LME results are compared between two separate analyses, with the first using data from only the first three time points (4, 10, and 16 months) and the second using data from all four time points (4, 10, 16, and 20 months), as is presented elsewhere in the manuscript. Brain-size-corrected volumes were modelled using an age by sex interaction with a random effect per subject. Age was expressed using a quadratic polynomial function to allow for non-linear change with age. The linear and quadratic components of age are written as poly(age,2)1 or poly(age,2)2, respectively. Effects of sex (sexF) were evaluated in females relative to males, along with the interaction of age and sex (poly(age,2)1:sexF and poly(age,2)2:sexF). Beta represents the standardized beta or coefficient value for each model term, with the adjacent column, Std.Error indicating the error associated with the standard beta measurement. The t-value for each model term is also included. Significant q-values, after 5% FDR correction, are denoted by blue shaded boxes, while those between 5 and 10% are shown by shaded grey boxes. L and R indicate a structure that is split over the left and right hemispheres, and was therefore reported and analyzed as two separate volumes.

**Supplementary Figures 1-8: Longitudinal trajectory of brain volumes with age and split by sex, for 71 unique brain regions in the aging Fischer rat.** MRI data were acquired longitudinally in rats aged 4-, 10-, 16- and 20-months old. Main effects of age and sex, as well as age by sex interactions, were determined using linear mixed effects modelling. Each rat is depicted by a single data point. The mixed effects model used to fit the data is represented by a line of best fit and 95% interval (shaded), split by sex. Data corresponding to males is shown using red circles with a solid line of best fit, while females are shown using blue triangles and a dashed line of best fit. Significance symbols are shown for each term in the model, where #^1^ and #^2^ represent the linear and second order age components of the natural spline with age, * denotes a main effect of sex in females relative to males, and †^1^ and †^2^ represent age by sex interactions with a linear or second order component of the natural spline with age, respectively. Multiple comparisons were corrected for using a 5% false discovery rate.

**Supplementary Figures 9 and 10: A side by side comparison between whole-brain voxelwise changes and regional changes as a result of age, sex, and their interaction, overlaid on the average anatomy background.** Brain-size-corrected volumes and relative Jacobians at each voxel were both modelled using an age by sex interaction with a random effect per subject. Age was expressed using a quadratic polynomial function to allow for non-linear change with age. The linear and quadratic components of the age term are written as poly(age,2)1 and poly(age,2)2, respectively. Effects of sex (sexF) were evaluated in females relative to males, along with the interaction of age and sex (poly(age,2)1:sexF and poly(age,2)2:sexF). The t-values for each term in the mixed effects model at either the voxelwise or structural level are plotted over the average anatomy background. t-value maps for the linear and quadratic age terms are shown in Figure 9, while those for the main effect of sex, and the interaction of age and sex are shown in Figure 10. The plot range for each set of t-values is specific to each model term and displays effects significant between 5 and 1% FDR, with t-values above 1% thresholded at the 1% value. The only exception to this is the t-statistic map for brain volumes under the poly(age,2)2:sexF heading, which are displayed with a 10% FDR cut-off. The scaled log relative Jacobian of peak voxels in the hindbrain, cerebellar lobule 3, and the hippocampus were visualized alongside the volume plots of the same structures, to highlight findings identified only at the voxel-wise level. A voxel from the temporal-parietal cortex is also visualized, alongside the volume data, to demonstrate findings only visible at the whole-volume level. For voxel and volume plots, the mixed effects model used to fit the data is represented by a line of best fit and 95% interval (shaded), split by sex. Data corresponding to males is shown using red circles with a solid line of best fit, while females are shown using blue triangles and a dashed line of best fit.

**Supplementary Figure 11: Visualization of age and sex effects, along with their interaction, on brain volumes (LME results) in healthy aging Fischer rats, thresholded to display the full range of t-values.** In the mixed effects model, volumes were each predicted by an age by sex interaction with a random intercept for each subject. Age was modelled using a quadratic polynomial function to account for the possibility of non-linear changes with age. The linear and quadratic components of age are written as poly(age,2)1 or poly(age,2)2, respectively. Effects of sex (sexF) were evaluated in females relative to males, and the interaction between age and sex was also examined (poly(age,2)1:sexF and poly(age,2)2:sexF, with males as the reference group. The regional t-values for each term in the mixed effects model are each shown in columns 1 through 5. The plot range for each set of t-values is specific to each model term and displays effects that range from significance at 5% FDR to the highest t-value except for the poly(age,2)2:sexF column which displays values significant at 10% FDR.

