## Supplementary Images 1-11 for "Longitudinal characterization of neuroanatomical changes in the Fischer 344 rat brain during normal aging and between sexes"

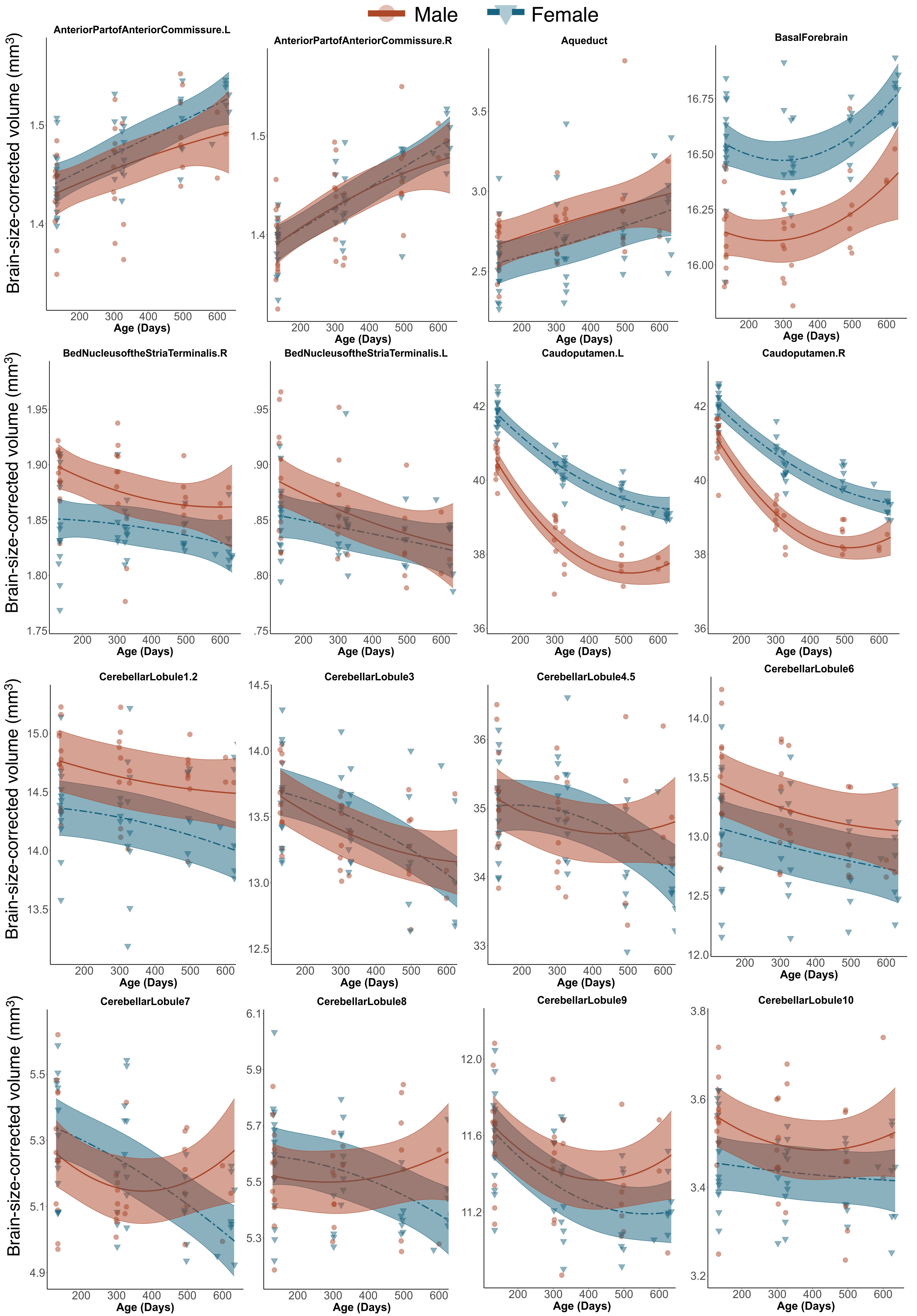

Male
Female
AnteriorPartofAnteriorCommissure.L
AnteriorPartofAnteriorCommissure.R
Aqueduct
BasalForebrain
BedNucleusoftheStriaTerminalis.R
BedNucleusoftheStriaTerminalis.L
Caudoputamen.L
Caudoputamen.R
CerebellarLobule1.2
CerebellarLobule3
CerebellarLobule4.5
CerebellarLobule6
CerebellarLobule7
CerebellarLobule8
CerebellarLobule9
CerebellarLobule10
Brain-size-corrected volume (mm³)
Age (Days)

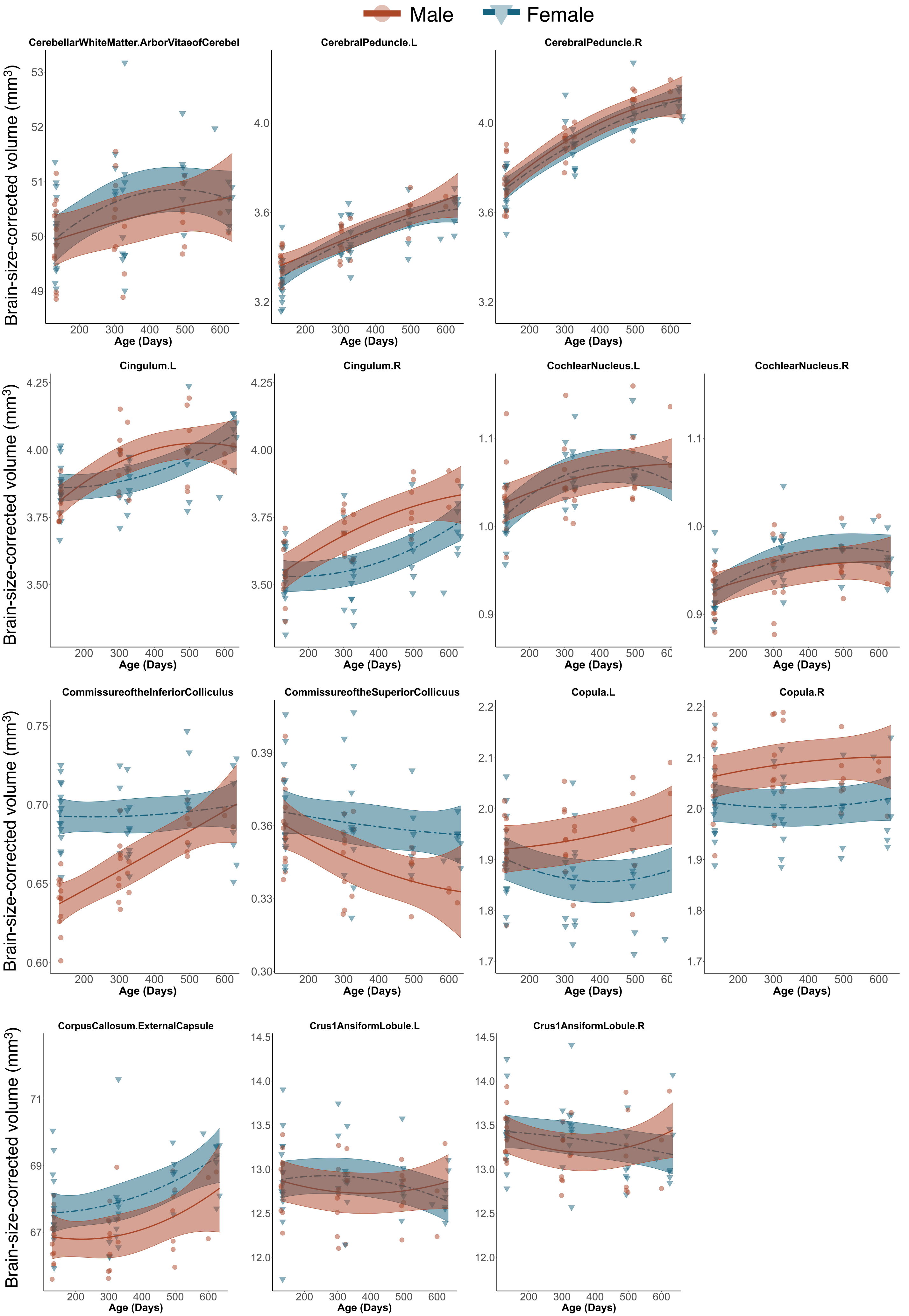

Male
Female
CerebellarWhiteMatter.ArborVitaeofCerebel
CerebralPeduncle.L
CerebralPeduncle.R
Cingulum.L
Cingulum.R
CochlearNucleus.L
CochlearNucleus.R
CommissureoftheInferiorColliculus
CommissureoftheSuperiorCollicuus
Copula.L
Copula.R
CorpusCallosum.ExternalCapsule
Crus1AnsiformLobule.L
Crus1AnsiformLobule.R
Brain-size-corrected volume (mm³)
Age (Days)

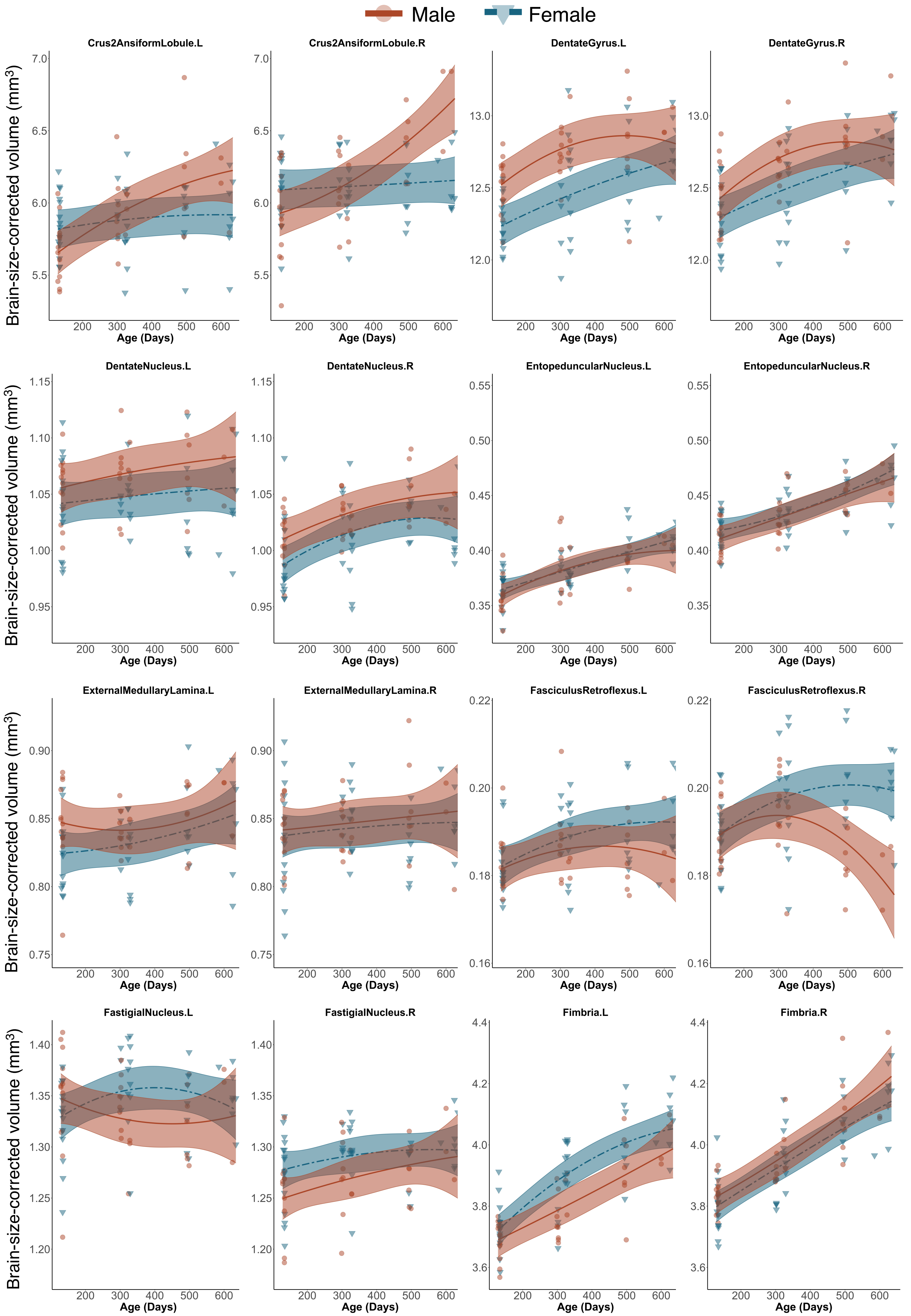

Male
Female
Crus2AnsiformLobule.L
Crus2AnsiformLobule.R
DentateGyrus.L
DentateGyrus.R
DentateNucleus.L
DentateNucleus.R
EntopeduncularNucleus.L
EntopeduncularNucleus.R
ExternalMedullaryLamina.L
ExternalMedullaryLamina.R
FasciculusRetroflexus.L
FasciculusRetroflexus.R
FastigialNucleus.L
FastigialNucleus.R
Fimbria.L
Fimbria.R
Brain-size-corrected volume (mm³)
Age (Days)

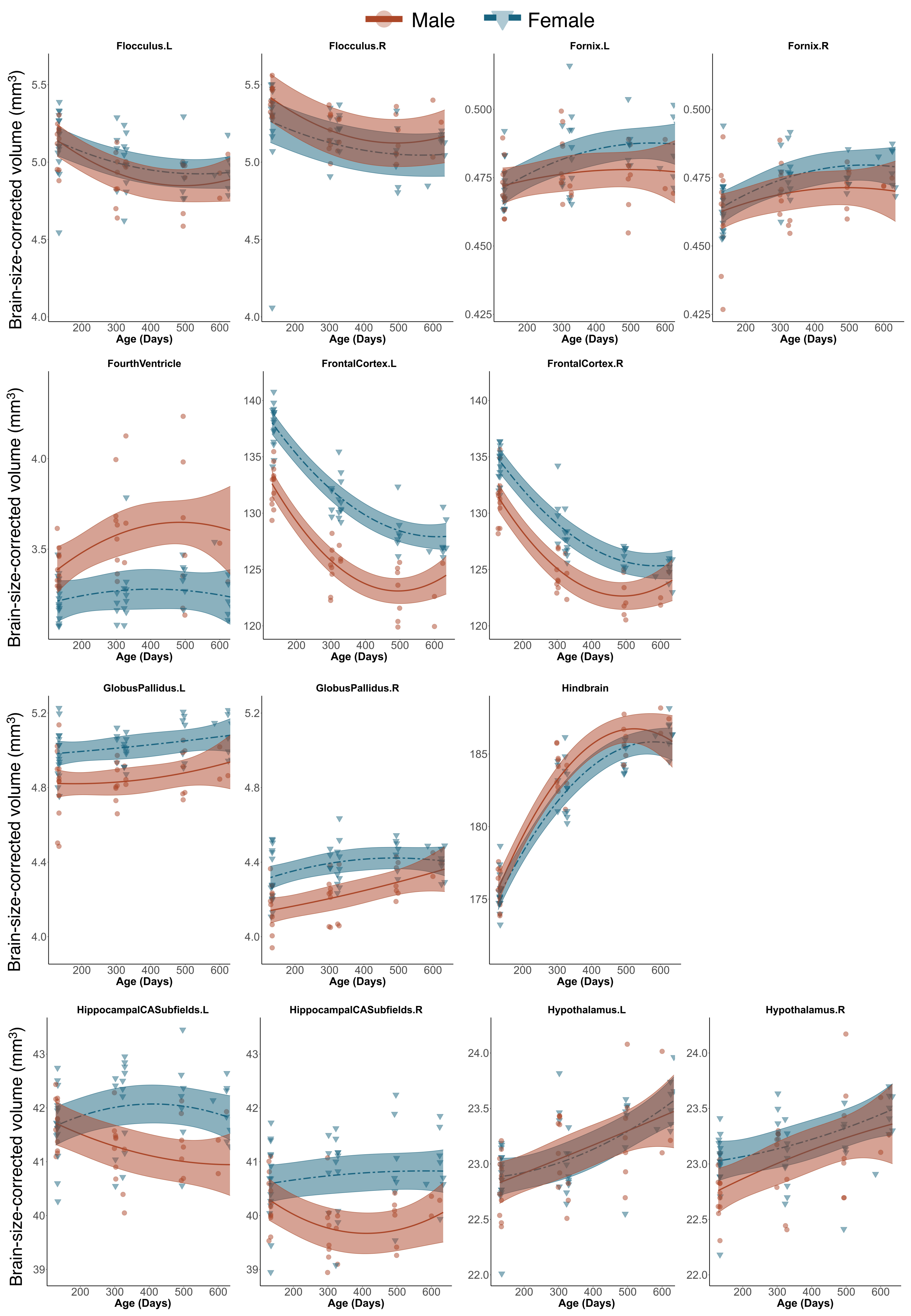

Male
Female
Flocculus.L
Flocculus.R
Fornix.L
Fornix.R
FourthVentricle
FrontalCortex.L
FrontalCortex.R
GlobusPallidus.L
GlobusPallidus.R
Hindbrain
HippocampalCASubfields.L
HippocampalCASubfields.R
Hypothalamus.L
Hypothalamus.R
Brain-size-corrected volume (mm³)
Age (Days)

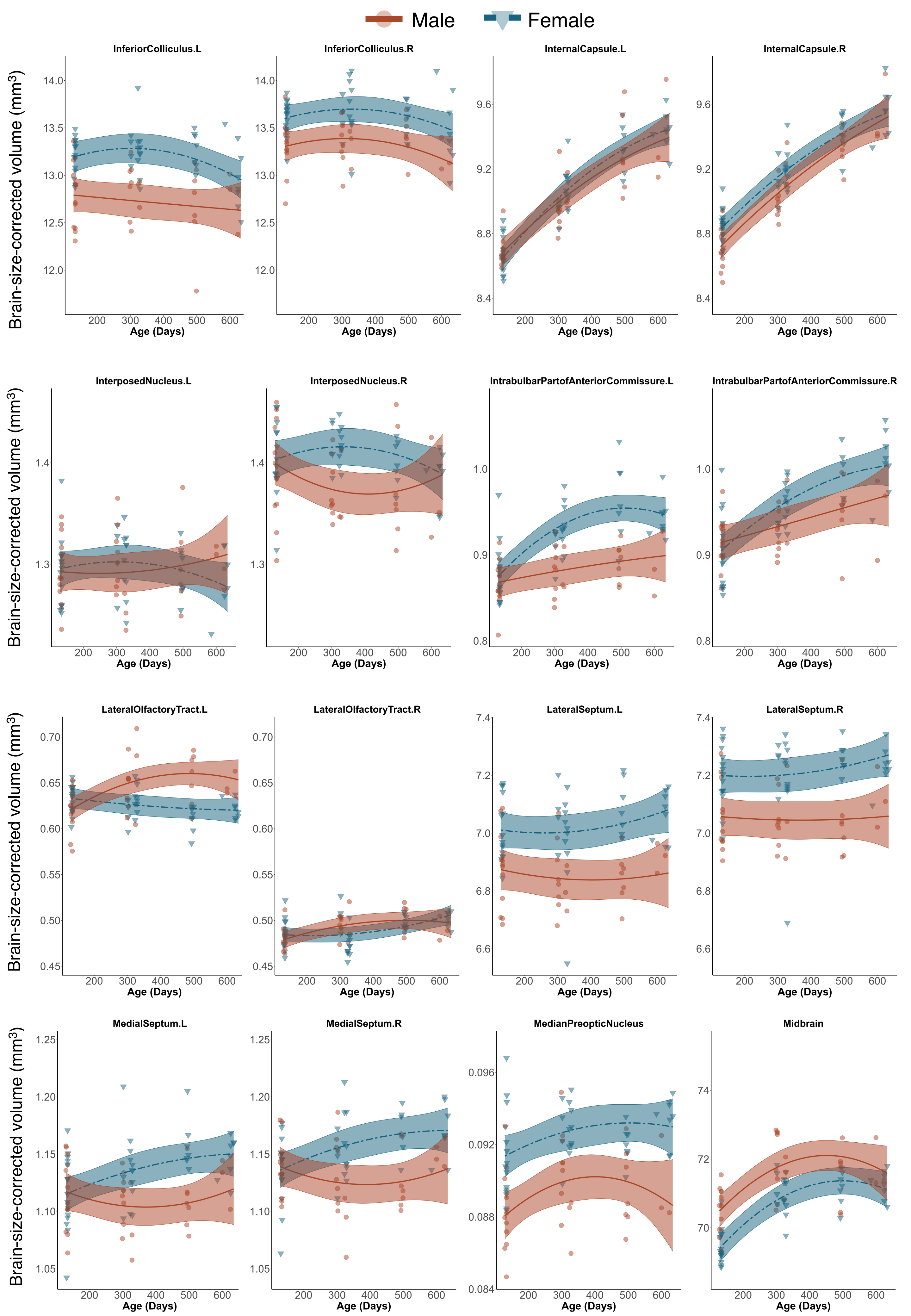

Male
Female
Brain-size-corrected volume (mm³)
Age (Days)
InferiorColliculus.L
InferiorColliculus.R
InternalCapsule.L
InternalCapsule.R
InterposedNucleus.L
InterposedNucleus.R
IntrabulbarPartofAnteriorCommissure.L
IntrabulbarPartofAnteriorCommissure.R
LateralOlfactoryTract.L
LateralOlfactoryTract.R
LateralSeptum.L
LateralSeptum.R
MedialSeptum.L
MedialSeptum.R
MedianPreopticNucleus
Midbrain

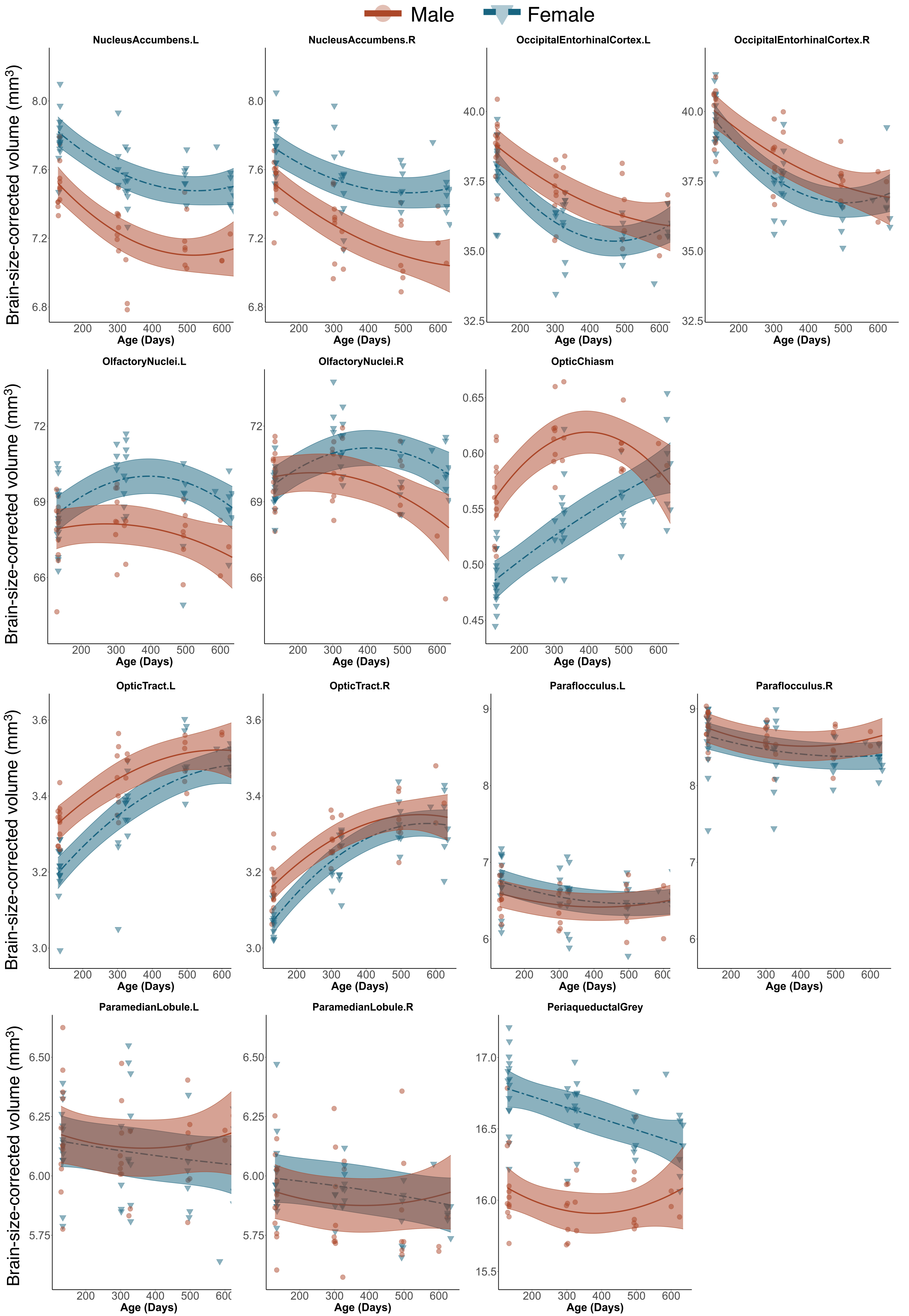

Male
Female
NucleusAccumbens.L
NucleusAccumbens.R
OccipitalEntorhinalCortex.L
OccipitalEntorhinalCortex.R
OlfactoryNuclei.L
OlfactoryNuclei.R
OpticChiasm
OpticTract.L
OpticTract.R
Paraflocculus.L
Paraflocculus.R
ParamedianLobule.L
ParamedianLobule.R
PeriaqueductalGrey
Brain-size-corrected volume (mm³)
Age (Days)

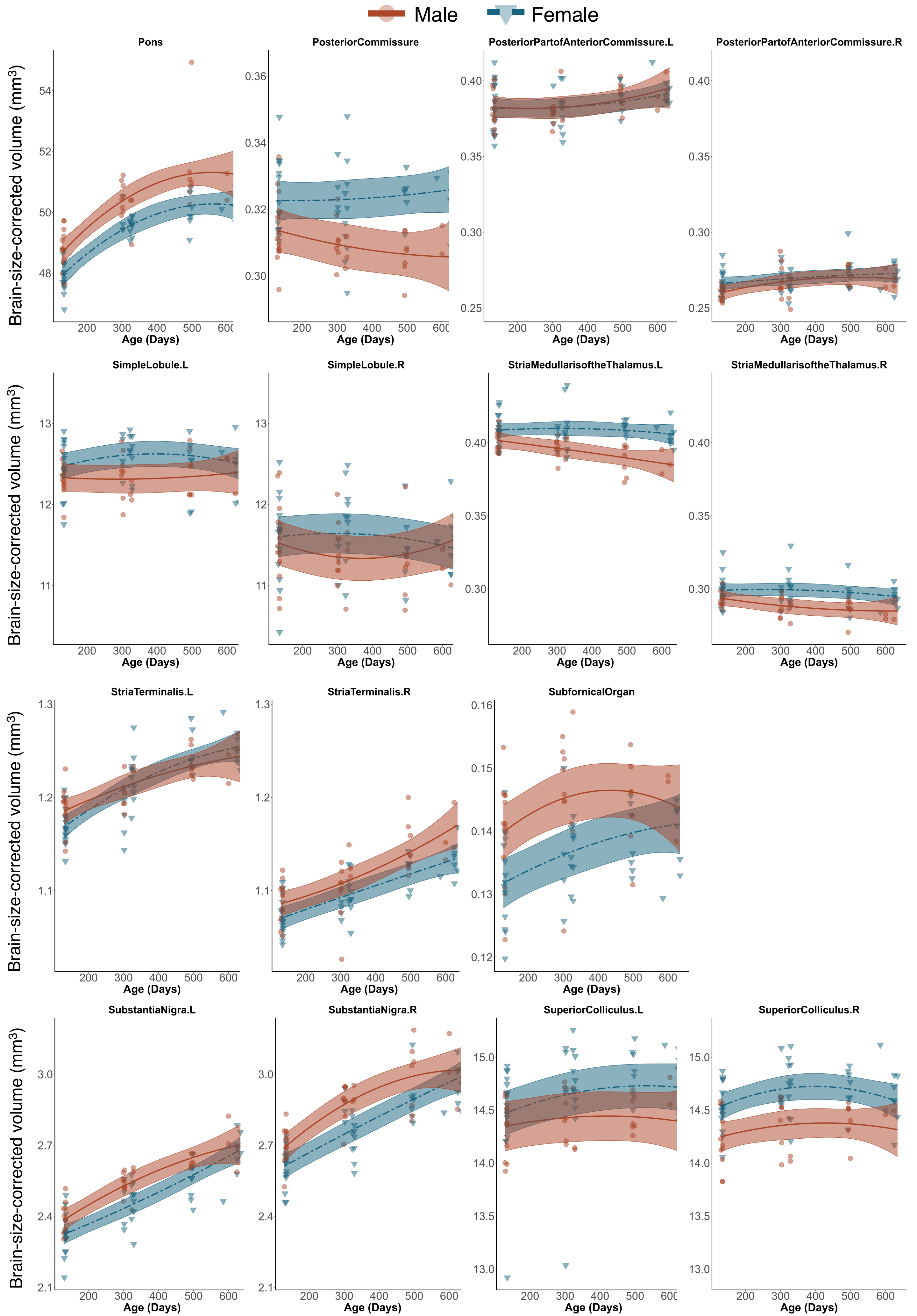

Male
Female
Pons
PosteriorCommissure
PosteriorPartofAnteriorCommissure.L
PosteriorPartofAnteriorCommissure.R
SimpleLobule.L
SimpleLobule.R
StriaMedullarisoftheThalamus.L
StriaMedullarisoftheThalamus.R
StriaTerminalis.L
StriaTerminalis.R
SubfornicalOrgan
SubstantiaNigra.L
SubstantiaNigra.R
SuperiorColliculus.L
SuperiorColliculus.R
Brain-size-corrected volume (mm³)
Age (Days)

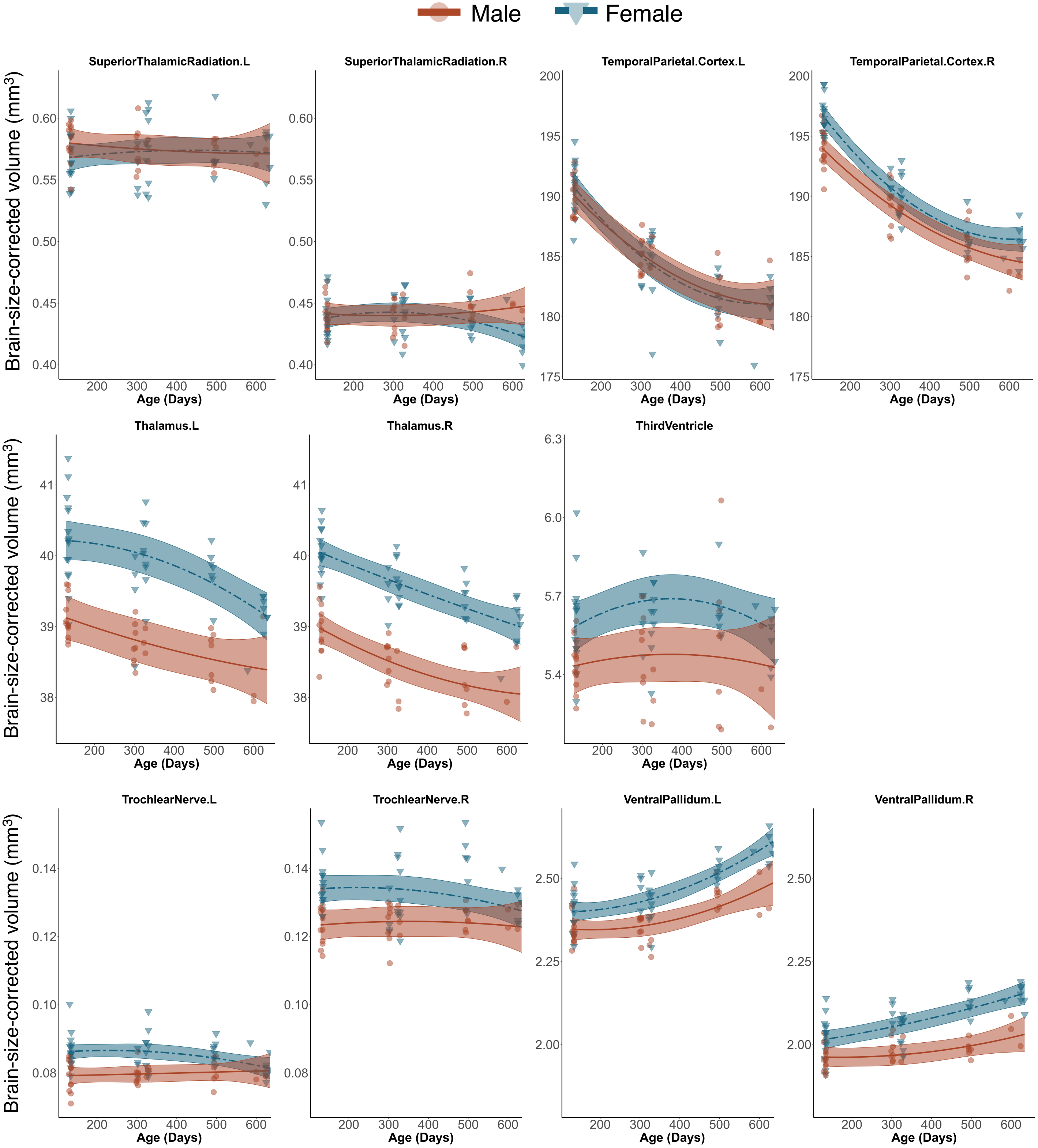

Male
Female
SuperiorThalamicRadiation.L
SuperiorThalamicRadiation.R
TemporalParietal.Cortex.L
TemporalParietal.Cortex.R
Thalamus.L
Thalamus.R
ThirdVentricle
TrochlearNerve.L
TrochlearNerve.R
VentralPallidum.L
VentralPallidum.R
Brain-size-corrected volume (mm³)
Age (Days)

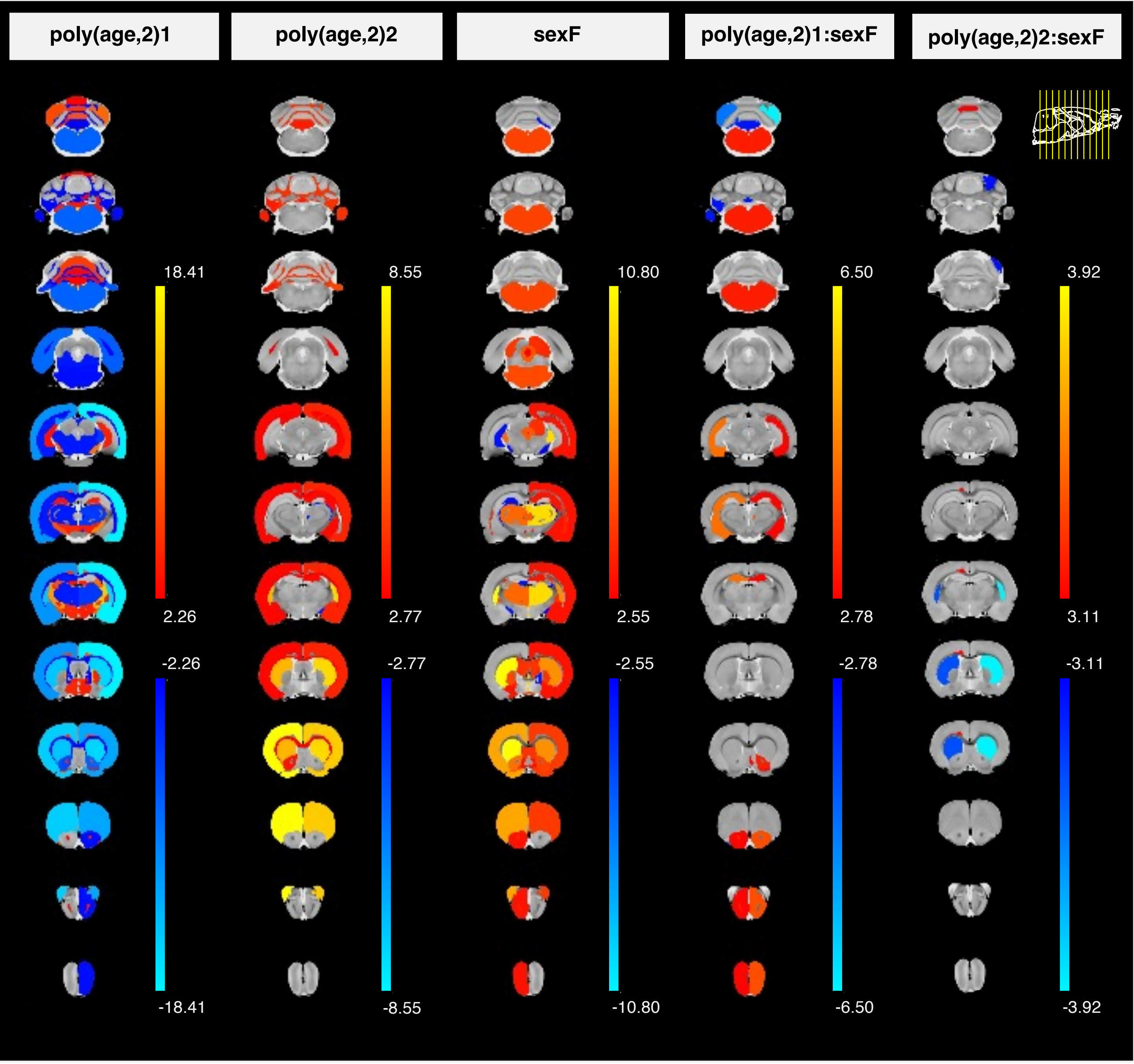

poly(age,2)1
poly(age,2)2
sexF
poly(age,2)1:sexF
poly(age,2)2:sexF
18.41
2.26
-2.26
-18.41
8.55
2.77
-2.77
-8.55
10.80
2.55
-2.55
-10.80
6.50
2.78
-2.78
-6.50
3.92
3.11
-3.11
-3.92

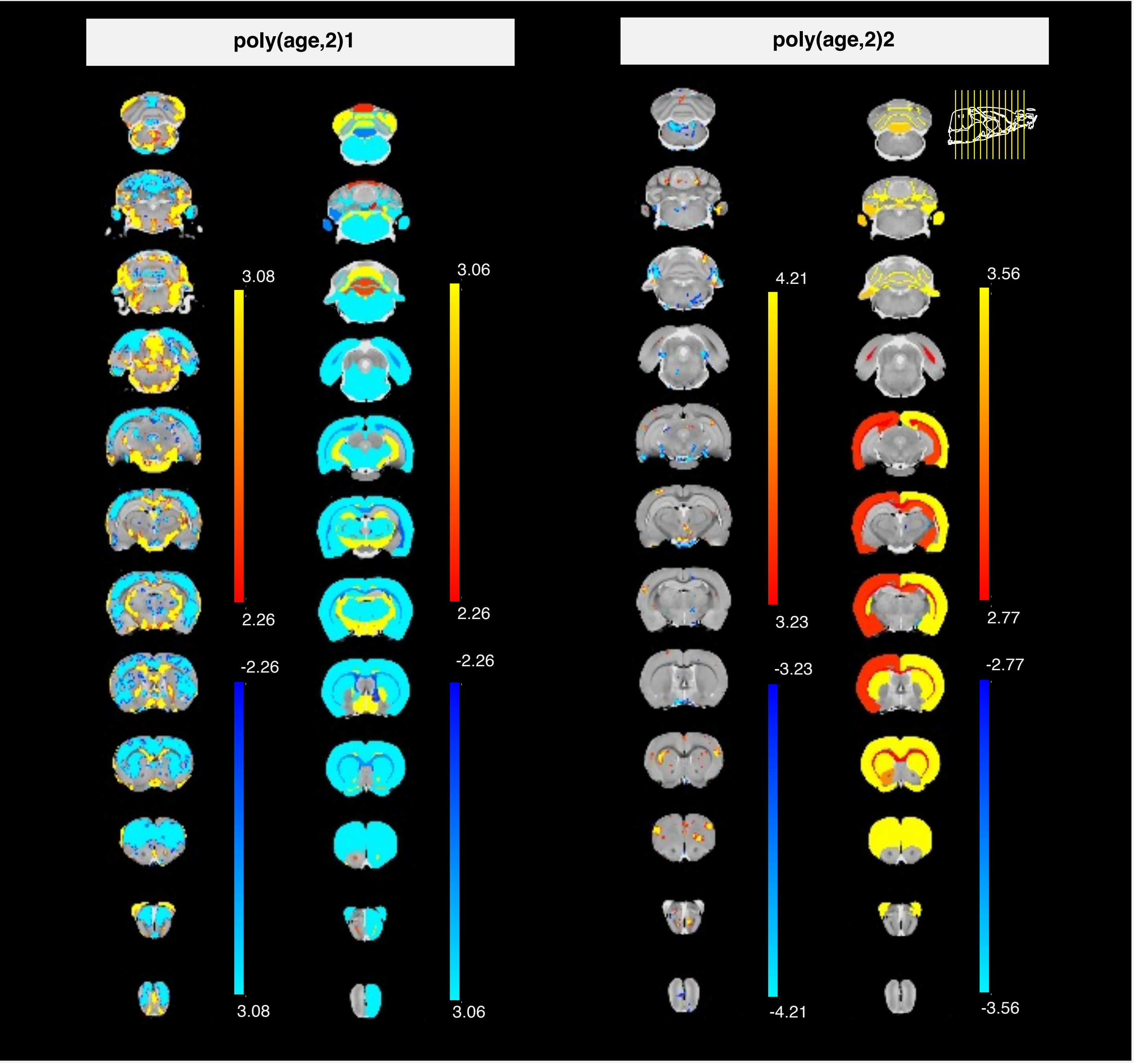

poly(age,2)1
poly(age,2)2
3.08
2.26
-2.26
3.08
3.06
2.26
-2.26
3.06
4.21
3.23
-3.23
-4.21
3.56
2.77
-2.77
-3.56

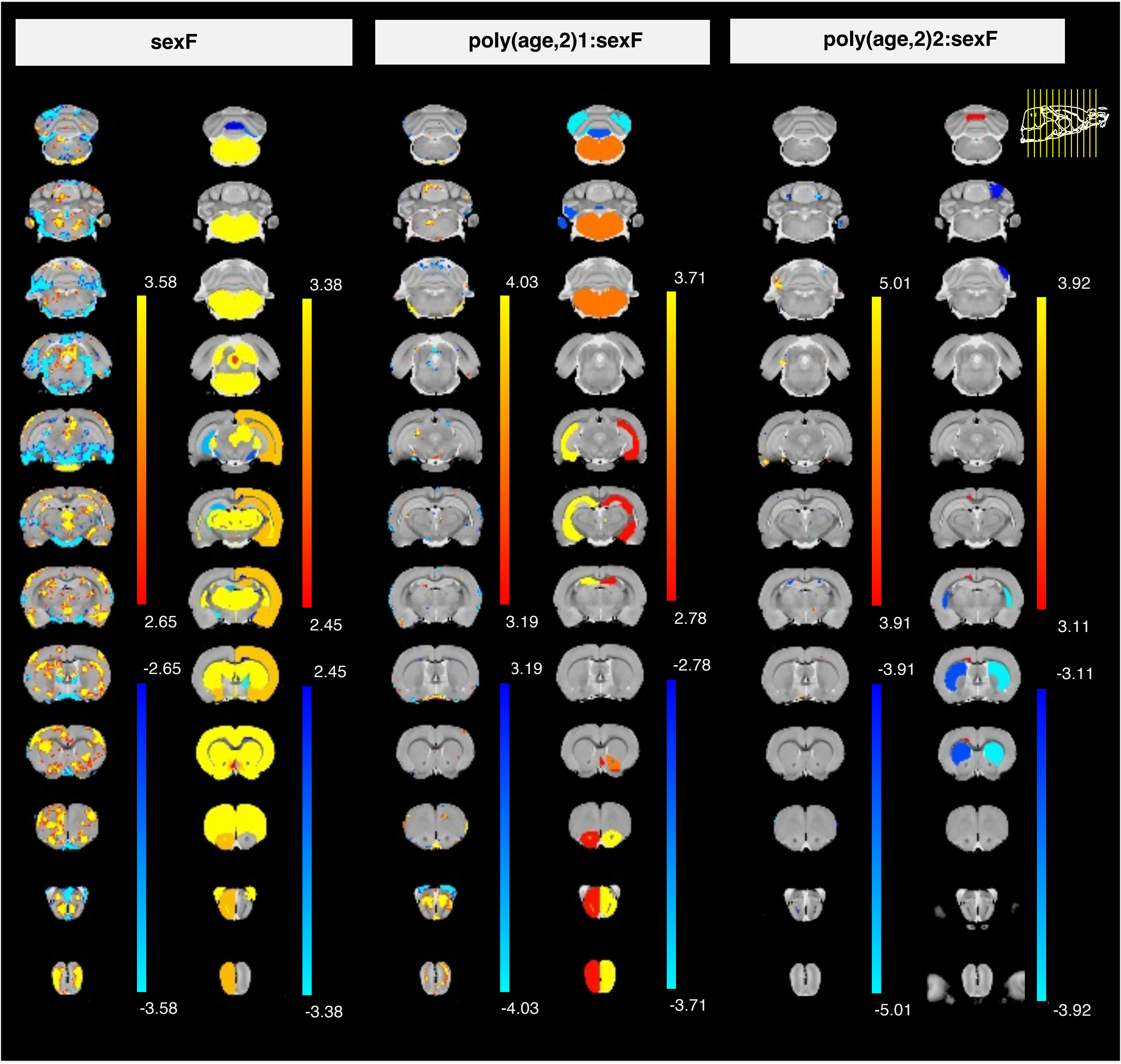

sexF
poly(age,2)1:sexF
poly(age,2)2:sexF
3.58
2.65
-2.65
-3.58
3.38
2.45
2.45
-3.38
4.03
3.19
3.19
-4.03
3.71
2.78
-2.78
-3.71
5.01
3.91
-3.91
-5.01
3.92
3.11
-3.11
-3.92
